# Electrostatic Engineering of Phosphoketolase Enhances Activity on Small Non-phosphorylated Sugars and Improves Cell-Free ATP Regeneration from Inexpensive C_2_-Substrates

**DOI:** 10.64898/2026.03.04.709640

**Authors:** Franziska Kraußer, Christopher M. Topham, Kenny Rabe, Jan-Ole Kundoch, Daniel Ohde, Andreas Liese, Thomas Walther

**Affiliations:** Chair of Bioprocess Engineering, Institute of Natural Materials Technology, TU Dresden, Bergstraße 120, 01062 Dresden, Germany; Molecular Forces Consulting, 24 Avenue Jacques Besse, 81500 Lavaur, France; Institute of Technical Biocatalysis, Hamburg University of Technology, Denickestr. 15, 21073 Hamburg, Germany

**Keywords:** Phosphoketolase, Acetyl phosphate synthesis, ATP cofactor regeneration, Molecular modelling, Multiple sequence alignment analysis, Protein residue pK_a_ prediction, electrostatic enzyme design and engineering

## Abstract

Phosphoketolases can be used to convert non-phosphorylated sugars to the high energy compound acetyl phosphate and the versatile metabolic precursor acetyl-CoA. The performance of these pathways is limited by low catalytic activity of natural phosphoketolases towards these sugars. Here, we report the rational engineering of the phosphoketolase from *Bifidobacterium adolescentis* (Bad.F6Pkt) to enhance its activity and affinity towards glycoaldehyde (GA) and D-erythrulose (ERU) through re-organisation of the protein electric field to reproduce the role of terminal phosphate groups in cognate substrates. Guided by predicted induced side-chain pK_a_ shifts, visualisation of electrostatic potential difference maps alongside molecular modelling and sequence variation analyses, we identified mutations that could promote *in situ* ring opening of the pre-dominant cyclic GA dimer form in solution. This approach to the electrostatic inverse design problem yielded the GA-specific double mutant H142N:E153D, exhibiting a ten-fold improved affinity and slightly enhanced catalytic efficiency (K_M_ = 4.4 mM, k_cat_/ K_M_ = 26.3 s^-1^ M^-1^) compared to the previously reported H142N variant (K_M_ = 42.3 mM, k_cat_/ K_M_ = 20.6 s^-1^ M^-1^). We additionally constructed a H256Y:H260Y:H548Y variant comprising long-range electrostatic mutations with a 3.8-fold increased catalytic efficiency (k_cat_/ K_M_ = 49.6 s^-1^ M^-1^) on the acylic four-carbon ERU ketose compared to the wild-type enzyme. The engineered enzymes were evaluated in cell-free enzyme cascades for ATP regeneration via acetyl phosphate formation. The H142N variant enabled efficient ATP regeneration from GA and ethylene glycol, whereas the H142N:E153D mutant exhibited reduced stability under synthesis conditions. Furthermore, coupling of a highly GA-specific D-threose aldolase and a D-threose isomerase with the PKT triple mutant enabled rapid conversion of GA into C_4_ sugar intermediates and significantly improved ATP regeneration from GA.

## Introduction

The physiological function of phosphoketolases is to cleave the phosphorylated sugars xylulose 5-phosphate (Xu5P) or fructose 6-phosphate (F6P) into acetyl phosphate (acetyl-P) and glyceraldehyde or erythrose-4-phosphate, respectively. Xu5P-specific PKTs are implicated in a xylose-degrading phosphoketolase pathway which is found in heterofermentative lactic acid bacteria. PKTs with activity on F6P facilitate degradation of hexoses in *Bifidobacteria* through the so called ‘bifid shunt’ ^1–4^. Acetyl-P can be readily converted to the metabolic hub molecule acetyl-CoA by the action of phosphate acetyltransferase (Pta)^5^. Phosphoketolases have consequently attracted considerable attention in metabolic engineering endeavours to implement synthetic metabolic pathways for the carbon-conserving production of acetyl-CoA ^6–8^.

Extending this concept, it has been recently shown that PKT enzymes are able to act on non-phosphorylated sugars, such as fructose, erythrulose and glycolaldehyde ^6,7,9^. This opens up the highly attractive prospect of applications that produce acetyl-P and downstream acetyl-CoA without the need to activate these sugars in ATP-consuming reactions. For instance, Lu et al. ^7^ have used PKTs to convert glycolaldehyde to acetyl-P. This enzyme-catalysed reaction is a crucial step in the implementation of linear synthetic pathways that convert methanol into acetyl-CoA without carbon loss ^10^. Kraußer et al have reported that enzymatic cascades can be designed that chain substrate specific PKT activities on D-fructose, D-erythrulose (ERU) and glycolaldehyde (GA) to produce three mol of acetyl-P from one mol of fructose ^9^. The high energy metabolite acetyl-P can be subsequently used to produce ATP via the acetate kinase reaction, thus, providing a convenient means to regenerate ATP from inexpensive sugars in cell-free reaction systems ^11^.

Although PKT applications that exploit non-phosphorylated substrates are of considerable commercial interest in biotechnology, cognate PKTs unfortunately have inherently poor catalytic activities on non-natural substrates. Recent enzyme engineering studies ^6,7,9^ using the template PKT enzyme from *Bifidobacterium adolescentis* (Bad.F6Pkt), which has high cognate substrate activity and can be produced at comparatively high levels in *Escherichia coli* ^9^, have focussed on improving catalytic performance towards non-phosphorylated ketose sugars and GA. We have obtained significant improvements in D-fructose, ERU and GA activity by single-mutation library screening at Bad.F6Pkt residue positions contacting a modelled bound open-chain covalent adduct reaction intermediate formed between the thiamine phosphate (TPP) cofactor and fructose-6-phosphate (F6P)^9^. Nevertheless, a growing body of evidence concerning the non-additive effects of single mutations on phosphoketolase^6,9^ and closely related transketolase^12^ activities on non-phosphorylated substrates strongly suggests that this type of epistasis is primarily electrostatic in origin. The coupling of the protein election field to an enzyme reaction is widely believed to play a determinant role in catalysis^13–15^. Combinatorially antagonistic mutation-induced perturbations of the protein electric field are thus likely to account for difficulties in recovering native-like PKT catalytic activities on non-phosphorylated substrates. Physics-based modelling methods originally developed for studying enzyme-catalysed reactions are increasingly being adapted to the diverse goals of computational enzyme design^16–20^. Improvements in enzyme variant catalytic properties through computational optimisation of local electric fields underpin recasting of protein sequence space exploration as the electrostatic inverse folding problem^13,21–24^.

In the present study we aimed at further improvement in the catalytic activity of Bad.F6Pkt on GA since the aldehyde is known to inactivate phosphoketolases and is toxic to cells when present at elevated concentrations ^10,25^. In parallel, we sought PKT catalytic efficiency improvements towards the C_4_ ketose ERU as GA can be conveniently converted to this sugar by chaining threose aldolase and threose isomerase reactions, thereby providing a means of maintaining low ambient GA concentration in the synthetic reaction pathway.

We have applied physics-based modelling techniques in conjunction with PKT family sequence variational analysis to fine tune the catalytic field in Bad.F6Pkt and induce side-chain pK_a_ shifts in key histidine base catalytic residues that appear to compensate for the absence of substrate terminal phosphate groups. Rational engineering of better adapted electrostatic pre-organisation in Bad.F6Pkt enabled the identification of a GA-specific H142N:E153D double mutant with a suitably operationally lowered K_M_ value of 4.4 mM. We additionally obtained an H256Y:H260Y:H548Y triple variant carrying co-operative long-range electrostatic mutations with a 3.8-fold increased catalytic efficiency towards ERU compared to the wild-type enzyme.

The newly constructed enzymes were employed in a cell free-reaction system to regenerate ATP from inexpensive ethylene glycol (EG) by chaining EG dehydrogenase, GA-dependent PKT and acetate kinase reactions (Figure 1). Additionally, we implemented a pathway to convert GA to ERU by coupling a highly GA-specific D-threose aldolase enzyme (*E. coli* fructose 6-phosphate aldolase variant Ec.FsaA L107Y:A129G^26^) with a D-threose isomerase (Figure 1). We showed that ATP regeneration from GA and EG was possible with the recently reported Bad.F6Pkt H142N variant^7^ providing the highest ATP yields. Conversion of GA to ERU via threose aldolase and threose isomerase reactions improved ATP regeneration with the highest yield obtained using the Bad.F6Pkt H256Y:H260Y:H548Y enzyme developed here.

**Figure 1.**
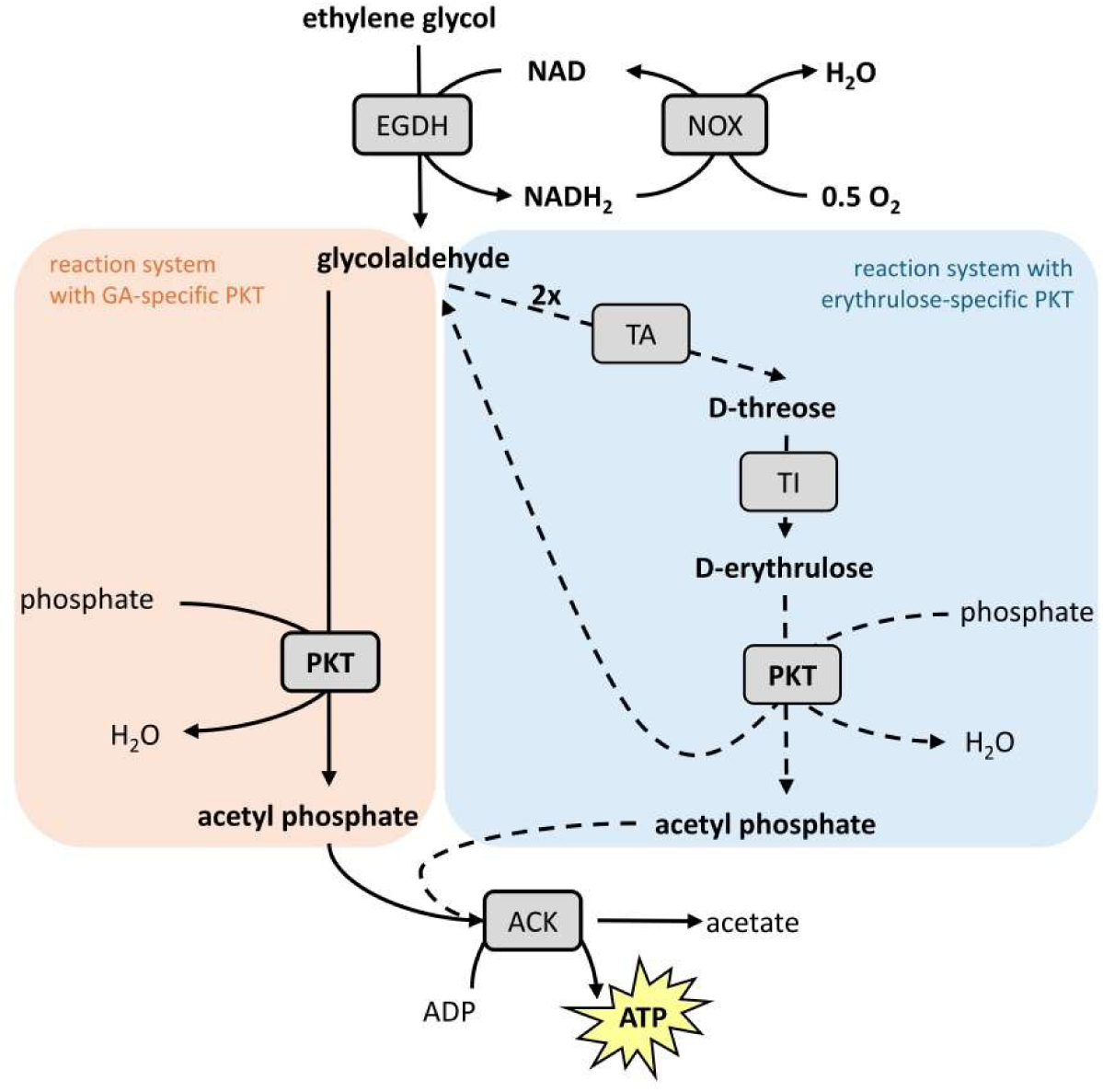
Enzymatic cascades for adenosine triphosphate (ATP) regeneration from ethylene glycol. Abbreviations: EGDH – ethylene glycol dehydrogenase, NOX – NADH oxidase, GA – glycolaldehyde, TA – threose aldolase, TI – threose isomerase, PKT – phosphoketolase, ACK – acetate kinase.

## Results

### Optimisation of the catalytic field in Bad.F6Pkt to promote catalysis on non-phosphorylated substrates

The phosphoketolase (PKT) enzyme from *Bifidobacterium adolescentis* (Bad.F6Pkt) was used here as the template for enzyme engineering of enhanced binding and activity towards ERU and GA non-phosphorylated substrates. Bad.F6Pkt is a member of the XFPK (EC 4.1.2.22) phosphoketolase sub-family present in *Bifidobacterium* species operating a bifid-shunt fermentation pathway. XFPK enzymes display comparable specificities for cyclic fructose-6-phosphate (F6P) and acyclic xylulose-5-phosphate (X5P) substrates, whilst the common PKT form present in most organisms (XPK, EC 4.1.2.9) is mainly specific for X5P (Scheidig et al 2019). Previously we identified a Bad.F6Pkt variant (H548N) with 5.6-fold increased activity on the non-phosphorylated substrate D-fructose, a 2.2-fold increase in ERU activity and a 1.3-fold increase in GA activity compared to the wild-type enzyme ^9^. However, this mutation in the mobile QN (Q546-N549) loop ^27^ in the substrate binding channel (Figure 2A) did not lower K_M_ for ERU or GA (Table 2), although this is a prerequisite for the implementation of an *in vitro* reaction system that avoids GA accumulation to reduce enzyme inactivation (see Figure 1).

**Figure 2.**
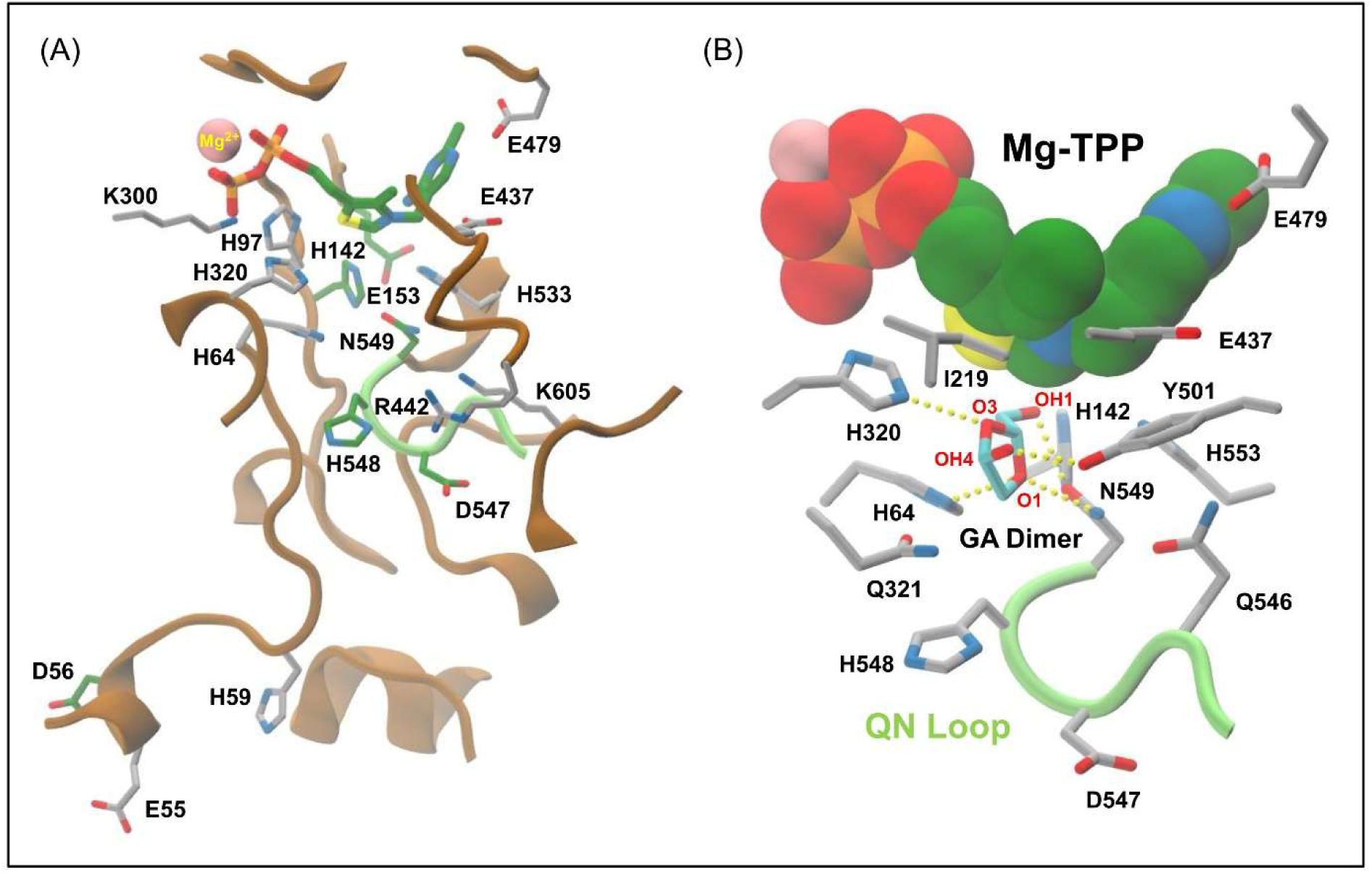
Substrate binding channel in Bad.F6Pkt. **(A)** The TPP stick representation is coloured according to element type: carbon, green; nitrogen, blue; oxygen, red; sulphur, yellow; phosphorus, orange. The bound Mg^2+^ ion is shown as a pink coloured van der Waals sphere. Sidechain carbon atoms at mutated enzyme residue positions are coloured in green, and in grey at ionisable residue positions used to monitor shifts in calculated pK_a_ values arising from mutation (see Methods). **(B)** The Mg-TPP cofactor is shown as van der Waals spheres using the same colour scheme. Bad.F6Pkt residue side-chain (grey carbons) good quality hydrogen bonding interactions with the cyclic GA dimer, 2-hydroxymethyl-4-hydroxy-1,3-dioxalane (Supplementary Information Figure S1) depicted with cyan coloured carbon atoms, are indicated by yellow solid sphere vectors.

The PKT reaction mechanism for the conversion of the cognate F6P and X5P substrates to acetyl-P is well established ^28–31^. Scheme S1 illustrates key reaction intermediates and catalytic steps for the multiple-step formation of acetyl-P from the acylic four-carbon ERU ketose. Computational simulation of the PKT-catalysed conversion of glycolaldehyde to acetyl-P ^7^ implicated the involvement of the reactive aldehyde group of the monomeric glycolaldehyde species and the participation of the protonated H553 residue tautomer in the formation of the α,β-dihydroxyethyl TPP (DHETPP) enzyme reaction intermediate ^28,31^. The alternative reaction pathway for PKT-catalysed GA conversion is integrated in Scheme S1, revealing catalytic steps in common with the reaction of ERU. GA exists in the solid crystalline state as a six-membered cyclic dimer (2,5-dihydroxy-1,4-dioxane) which dissociates in aqueous solution via an acyclic intermediate to yield a re-cyclised dimer, identified as 2-hydroxymethyl-4-hydroxy-1,3-dioxolane (Figure S1), along with different monomeric forms^32^.

Quantitative analysis indicates that GA is predominantly present in solution as a hydrated monomer (70-75%), with the five-membered 2-hydroxymethyl-4-hydroxy-1,3-dioxolane cyclic dimer accounting for approximately 17 %. A mere 4-5 % of GA exists in its monomeric form with a reactive aldehyde moiety^33,34^. The capacity of PKT to catalyse GA conversion would appear to be impaired by the low abundance of the reactive substrate species, acting as an effective brake on net flux through the synthetic reaction pathway for the generation of ATP.

Enzyme catalytic efficiency has been postulated by Warshel ^35,36^ to originate from the intra-molecular protein electric field that permanently favours the reaction transition-state (TS) over the reactants. Protein electrostatic pre-organisation not only provides favourable enthalpic contributions to TS binding, but also permits enzyme-catalysed reactions to avoid an entropic penalty associated with the re-organisation of the environment during TS crossing. Vibrational Stark effect (VSE) spectroscopy has been used to experimentally demonstrate the correlation between enzyme catalytic rate enhancement and the magnitude of the electric field in the active-site ^37^. The k_cat_/K_M_ enzyme specificity constant kinetic parameter corresponds to the rate constant for the reaction of free enzyme with substrate ^38^. According to transition-state theory, k_cat_/K_M_ can be related to the free energy difference between the rate limiting enzyme-reaction TS energy barrier and the free enzyme state. This conveniently allows changes in k_cat_/K_M_ arising from mutation or other perturbation to be inferred from predicted modulation of the protein electric field in the free or unbound enzyme state.

Here we employ the Poisson-Boltzmann (PB) continuum electrostatics model as a computationally tractable framework for the calculation of ionisable protein residue pK_a_ values, electrostatic potential (φ(**r**)) and field strength ^18,39–41^ in mutant and wild type enzyme complexes with the Mg-TPP cofactor. Although calculated mutant - wild-type difference electrostatic potential (Δφ(**r**)) grids cannot of course provide any direct information concerning mutant- and/or substrate-dependent rate-limiting reaction steps, this can sometimes be inferred in our rational approach to PKT re-design from experimentally determined kinetic parameter feedback obtained by probe mutation and induced shifts in predicted residue side-chain pK_a_ values. The PB-based model allows for electrostatic polarisation to be taken into account of at all positions in the Bad.F6Pkt-TPP dimer. This is particularly advantageous in the fine-tuning of local electric fields as a means of improving catalysis towards non-phosphorylated substrates in mutant enzymes.

Bad.F6Pkt is an acidic protein with an isoelectric point (pI) of 4.2, determined by simulated titration (Figure S2) using DelPhiPKa ^42,43^. The observed field line high density in the substrate binding channels of the dimeric enzyme is in direct proportion to the electric field strength (Figure S3). Of the five identifiable ionisable histidine residue positions in the wild-type active-site (Figure 2), four (64, 97, 142 and 320) are fully buried with side-chain relative solvent accessibility (RSA) values below 7%. The imidazole side-chain of H548 in the QN loop is partially buried, principally through interaction with the neighbouring D547 acidic residue, with an RSA value (29%) in range 7-40%. Calculated side-chain pK_a_ values of all the histidine residues in the active-site apart from H97 are lower than experimentally determined macroscopic side-chain pK_a_ values of 6.02 ^44^ and 6.20 ^45^ for L-histidine in aqueous solution (see Table 1).

**Table 1.** Computed residue pK_a_ values in Bad.F6Pkt and shifts due to PO_4_. ^2^**^-^ binding and mutation.** Calculations at ionisable residue positions were carried on modelled wild-type and mutant Bad.F6Pkt complexes with Mg-TPP using DelPhiPKa as described in Methods. Tabulated predicted pK_a_ values at selected positions are averages over both chains in the dimeric enzyme. For clarity, only Induced pK_a_ shifts relative to the wild type enzyme (ΔpK_a_) of ≥ ±0.1 in magnitude are indicated below in parentheses. RSA, side-chain residue solvent accessible surface area relative to that in corresponding extended conformation Ala-XXX-Ala model reference tri-peptide structure ^46^. ^(a)^ Bound PO_4_^2-^ ion atomic co-ordinates were abstracted as the terminal phosphate group of the open-chain form of F6P in an overlaid copy of the modelled TPP-F6P (T6F) covalent adduct complex with Bad.F6Pkt ^9^. NA, not applicable. Undet., could not be determined.

| Bad.F6Pkt<br>Residue<br>Position | RSA<br>(%) | Wild<br>Type | Wild<br>Type<br>+ PO<sub>4</sub><sup>2-</sup> (a) | H548N | H548Y | H260Y:<br>H548Y | H256Y:<br>H260Y:<br>H548Y | H142N | H142N:<br>E153D |
| --- | --- | --- | --- | --- | --- | --- | --- | --- | --- |
| D26 | 13 | 1.97 | 1.97 | 1.97 | 1.97 | 2.42<br>(+0.45) | 2.76<br>(+0.79) | 1.97 | 1.97 |
| H59 | 26 | 5.55 | 5.63 | 5.63 | 5.63 | 5.64 | 5.63 | 5.58 | 5.58 |
| H64 | 2 | 5.08 | 5.70<br>(+0.62) | 5.28<br>(+0.20) | 5.40<br>(+0.32) | 5.42<br>(+0.34) | 5.41<br>(+0.33) | 5.42<br>(+0.34) | 5.47<br>(+0.39) |
| H97 | 0 | 6.09 | 6.13 | 6.09 | 6.09 | 6.10 | 6.10 | 6.16 | 6.20<br>(+0.11) |
| H142 | 2 | 4.90 | 4.98 | 4.96 | 4.95 | 4.96 | 5.00<br>(+0.10) | NA | NA |
| E153 | 21 | 2.23 | 2.27 | 2.25 | 2.25 | 2.26 | 2.26 | 2.43<br>(+0.20) | 1.81<br>(-0.43) |
| H256 | 46 | 6.08 | 6.08 | 6.08 | 6.08 | 6.21<br>(+0.13) | NA | 6.08 | 6.08 |
| H260 | 0 | 6.38 | 6.38 | 6.38 | 6.39 | NA | NA | 6.41 | 6.42 |
| K300 | 0 | 11.86 | 12.02<br>(+0.16) | 11.85 | 11.86 | 11.86 | 11.86 | 11.86 | 11.87 |
| H320 | 2 | 4.64 | 5.18<br>(+0.54) | 4.70 | 4.74<br>(+0.10) | 4.75<br>(+0.11) | 4.76<br>(+0.12) | 4.66 | 4.67 |
| E437 | 1 | 2.23 | 2.47<br>(+0.23) | 2.28 | 2.28 | 2.28 | 2.28 | 2.30 | 2.30 |
| R442 | 31 | 13.16 | Undet. | 13.15 | 13.14 | 13.15 | 13.15 | 13.17 | 13.16 |
| E479 | 1 | 2.58 | 2.63 | 2.59 | 2.59 | 2.59 | 2.59 | 2.63 | 2.63 |
| D547 | 26 | 1.19 | 1.64<br>(+0.45) | 1.40<br>(+0.21) | 1.40<br>(+0.21) | 1.40<br>(+0.21) | 1.40<br>(+0.21) | 1.22 | 1.23 |
| H548 | 29 | 5.52 | 6.51<br>(+0.99) | NA | NA | NA | NA | 5.52 | 5.53 |
| H553 | 0 | 4.93 | 5.38<br>(+0.45) | 5.02 | 5.04<br>(+0.11) | 5.04<br>(+0.11) | 5.03<br>(+0.10) | 5.14<br>(+0.21) | 5.20<br>(+0.27) |
| K605 | 19 | 10.76 | 12.98<br>(+2.22) | 10.80 | 10.80 | 10.80 | 10.80 | 10.77 | 10.76 |

*In situ* ring-opening of cyclic substrates such as D-fructose and the GA dimer species is potentially rate-limiting in the PKT-catalysed conversions of these non-phosphorylated substrates to acetyl-P. The β-furanose ring form of the cognate PKT substrate fructose 6-phosphate is believed to be linearized in the active site either by protonation of the ring oxygen (acid catalysis) or deprotonation of the anomeric C2 hydroxyl group (base catalysis), both of which have been suggested to be assisted by the 6-phosphate group via mediating solvent molecules ^31^. Since 2-hydroxymethyl-4-hydroxy-1,3-dioxalane does not possess an anomeric hydroxyl group, ring opening of the GA cyclic dimer may be expected to occur by acid catalysis. An energy minimised model complex of wild type Bad.F6Pkt enzyme with Mg-TPP and the bound cyclic GA dimer is shown in Figure 2B. The C2 ring atom in the GA dimer is separated by approximately 4 Å from the C2 TPP thiazolium ring carbon with which a covalent adduct is formed in the first step of the cognate F6P substrate reaction mechanism ^28,31^. The O1 ring oxygen is hydrogen bonded by the flanking N549 and H64 side-chains, while N549 can make an additional hydrogen bond with the 2-hydroxymethyl group. Substrate-specific interactions of the open chain TPP-F6P covalent adduct with the N549 residue in the *Bifidobacterium breve* (XFPK) enzyme were reported in recent docking simulation studies ^47^. H320 and Y501 are respectively able to form hydrogen bonds with the GA-dimer O3 ring oxygen and the 4-hydroxyl group. H64 has been proposed as a putative proton acceptor in the deprotonation of the C3 hydroxyl in C3-C4 bond cleavage of the natural substrate F6P ^29,30^. Acid-catalysed GA dimer ring opening could be mediated by a solvent proton or the direct participation of the protonated H64 side-chain tautomeric state.

The five-membered 2-hydroxymethyl-4-hydroxy-1,3-dioxalane GA cyclic dimer (Figure S1) notably bears a close structural resemblance to the β-furanose ring form of F6P, whereas D-fructose is sterically constrained to predominantly exist in the six-membered β-pyranose ring form ^48^. The postulated involvement of the 6-phosphate group in F6P ring opening ^31^ prompted us to investigate the influence of a single bound PO_4_ ^2-^ ion, extracted from a TPP-F6P model complex ^9^, on the calculated side-chain pK_a_ values of ionisable residues lining the substrate binding channel (Table 1). The predicted pK_a_s of the catalytic reaction centre H64, H320, and H553 residue sidechains were respectively shifted upwards by the presence of the phosphate ion by 0.62, 0.54 and 0.45 units. The calculated pK_a_ of the D547 residue sidechain close to PO_4_ ^2-^ ion binding site was also raised to a similar degree (0.45). The H64 and H320 residues have been variously inferred from substrate-adduct docking simulations to be the B_2_ catalytic base (Scheme S1) in XFPK enzymes responsible for deprotonation of the F6P C3 hydroxyl group in the second (aldose phosphate product formation) reaction step ^31^, whereas the H553 equivalent residue (H559) in the *Lactococcus lactis* XPK enzyme was assigned as the B_2_ catalyst ^47^. The B_2_ catalyst is also involved in the following DHETPP dehydration step common to all substrate reaction mechanisms ^31^ as indicated in Scheme S1. It is noteworthy that in the docked and cross-docked simulated PKT enzyme-substrate complexes reported by Scheidig *et al*.^47^, the terminal phosphate group of the X5P-TPP covalent adduct was positioned closer to the three catalytic reaction centre histidine residues with predicted perturbed pK_a_ values incurred by PO_4_ ^2-^ ion binding, than was the 6-phosphate group of F6P-TPP.

Collectively these findings suggest that in addition to an established steric role in the preferential stabilisation of the β-furanose ring form of the cognate F6P substrate, the 6-phosphate group may not only assist ring opening in F6P by raising the pK_a_ of H64 in the catalytic reaction centre of XFPK enzymes but may also aid deprotonation of TPP-F6P adduct via modulation of the B_2_ catalytic base pK_a_. Electrostatic mutations in Bad.F6Pkt able to mimic the induced effects of the terminal phosphate groups in cognate substrates can thus be anticipated to promote ring opening of the prevalent GA-dimer form in solution, with concomitant *in situ* release of the reactive GA monomer ^7^, as well as facilitating deprotonation of the linear TPP-ERU covalent adduct. Other protein electric field modulating mutations targeting the DHETPP dehydration reaction step are expected to improve catalytic activity towards both phosphorylated and non-phosphorylated substrates.

Whilst saturation by the cyclic D-fructose non-phosphorylated substrate could not be experimentally observed in the Bad.F6Pkt H548N mutant with the previously reported highest initial reaction rate, k_cat_/K_M_ for the acyclic ERU substrate was increased by approximately 1.6 fold compared to the wild-type enzyme, indicative of improved binding of the enzyme-substrate reaction TS ^9^. H548 forms part of the binding site for the terminal phosphate group in the complex with the covalent F6P-TPP substrate adduct. The calculated respective upward shifts of +0.20 and +0.21 in the pK_a_ values the H64 and D547 residue sidechains in the H548N mutant (Table 1), although smaller, are comparable with corresponding shifts elicited upon introduction of the charged PO_4_ ^2-^ ion surrogate for the 6-phosphate group in F6P. Stabilisation of the D547 sidechain carboxylate group by the mutant asparagine 548 sidechain carbamoyl group was identified as being mainly responsible for the rise in the predicted pK_a_ of D547. The electrostatic potential difference (Δφ(**r**)) map for the H548N mutant reveals local charge polarisation in the enzyme reaction centre and in an increase in the electric field strength parallel to the axis of the covalent bond formed with the C2 TPP thiazolium ring carbon (Figure 3A).

**Figure 3.**
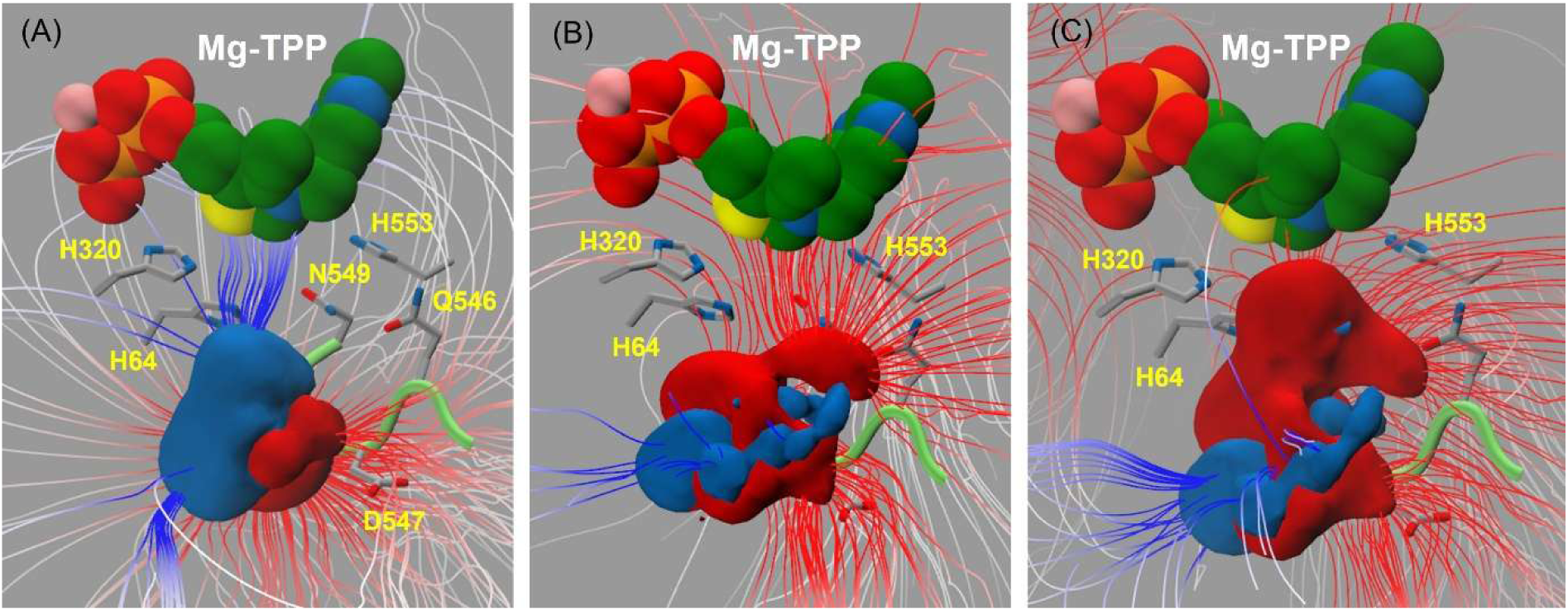
Electrostatic potential difference grid maps for Bad.F6Pkt variants carrying H548 mutation. **(A)** H548N **(B)** H548Y **(C)** H260Y:H548Y. Difference electrostatic potential (Δφ(**r**)), calculated relative to wild-type Bad.F6Pkt, is displayed as solid rendered ±2 kT/e contoured isosurfaces, coloured in blue or red for positive or negative Δφ(**r**), respectively. Electrostatic field lines, calculated at a fixed gradient magnitude setting, are coloured according to the electrostatic difference potential and point away from positive charge difference sources. The Mg-TPP cofactor is shown as van der Waals spheres with green coloured carbon atoms. Other atoms are coloured according to element type: nitrogen, blue; oxygen, red; sulphur, yellow; phosphorus, orange. The Mg^2+^ ion is shown in pink. Active-site histidine (64, 320 and 553) and QN loop (Q546, D547, H548, N549) residue side-chains are shown as in stick-form with grey-coloured carbons. The QN loop mainchain is depicted in cartoon form as a lime green tube.

### Electric field modulation as a design strategy to improve Bad.F6Pkt activity towards GA

We next sought to identify electrostatic mutations with the potential to facilitate GA dimer ring opening both at long range (Figure 4A) and at positions in the active-site centre and substrate binding channel of the Bad.F6Pkt enzyme (Figure 2A). Electrostatic re-organisation in mutants was evaluated from induced shifts in predicted pK_a_ values (ΔpK_a_) at ionisable (reporter) residue positions, including those most influenced by the introduction of a PO_4_ ^2-^ ion in the terminal F6P phosphate group binding site (Table 1), and by visual inspection of the corresponding (Δφ(**r**)) electrostatic potential difference maps and field line density at the active-site centre. Candidate replacement residue type selection was assisted by position-specific frequency profiles of tolerated residue substitution and statistical correlation energy measures of pairwise residue type dependence determined by analysis of a multiple sequence alignment of PKT enzyme family members (see Methods).

**Figure 4.**
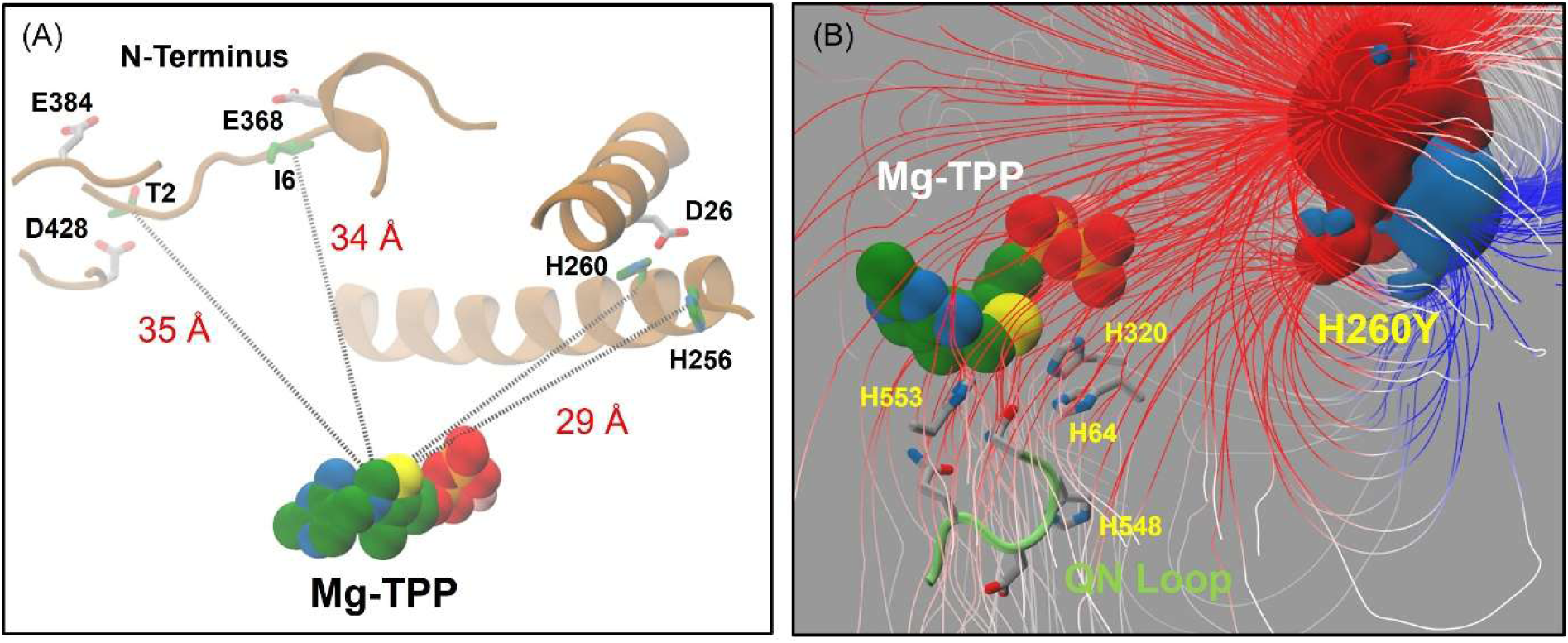
Long range electrostatic mutations in Bad.F6Pkt. **(A)** Locations of mutated residues (green side-chain carbons) and proximal acidic residues (grey side-chain carbons) with respect to the Mg-TPP cofactor in the enzyme catalytic centre. Mg-TPP is depicted with space filling van der Waals spheres coloured according to element type: carbon, green; nitrogen, blue; oxygen, red; sulphur, yellow; phosphorus, orange; magnesium, pink. Distance separations between the TPP C2 thiazolium ring atom and C^α^ atoms of mutated enzyme residues are annotated in red. **(B**) H260Y mutant electrostatic potential difference grid map. Solid rendered ±2 kT/e contoured isosurfaces are coloured in blue or red for positive or negative difference potential, respectively. Electrostatic field lines, calculated with the same gradient magnitude setting as in Figure 3, are coloured according to the electrostatic difference potential due to mutation. Histidine (64, 320 and 553) and QN loop (Q546, D547, H548, N549) residue side-chains in the active-site are shown as in stick-form with grey-coloured carbons. The QN loop mainchain is depicted in cartoon form as a lime green tube.

#### (1) Long-range mutations at T2, I6, H256 and H260 positions

Long-range H260Y and T2A:I6T Bad.F6Pkt mutations identified by random mutagenesis were reported by Dele-Osibanjo *et al*. ^49^ to increase k_cat_/K_M_ towards F6P. A further enhancement in catalytic efficiency could be obtained by combining mutations at the three sites in the same region close to the enzyme surface approximately 29 Å (H260Y) and 34 to 35 Å (T2A:I6T) removed from the active-site. Interestingly no changes in K_M_ were observed, indicating that the increases in catalytic efficiency derived directly from favourable increases in reaction TS binding rather than substrate ground-state binding. The exhibition of relatively larger mutant rate enhancements towards F6P as compared to the acyclic X5P substrate ^49^ is consistent with a lowering of the reaction energy barrier for ring opening, although differential effects on the deprotonation of the substrate TPP covalent adducts cannot be ruled out. Mutation induced changes to the protein electric field in the active-site are illustrated in Figure 4B (H260Y) and Figure S4 (T2A:I6T). In contrast to the H548N mutant (Figure 3A), charge polarisation due to mutation leads to an excess of negative electrostatic potential in the catalytic centre. Whereas catalytic field modulation arising from the long-range H260Y mutation is confined to the closest active-site centre in the Bad.F6Pkt dimer, perturbation of the electric field can be observed in both catalytic centres in the T2A:I6T long-range double mutant.

Analysis of predicted ΔpK_a_ data in long-range electrostatic mutations (Table S2) did not reveal any change in pK_a_ values of ionisable residue sidechains in the active-site. An increase of 0.45 in the predicted sidechain pK_a_ of D27 was observed in the H260Y mutant. The partially buried D27 residue position lies on an adjacent α-helix to H260 which shields the carboxylate side-chain from the solvent (Figure 4A). Although histidine is the most prevalent residue type at this position in the PKT family multiple sequence alignment (MSA), tyrosine is well-represented (Figure S6). A similar local environmental change accompanying tyrosine mutation of H256, located at the N-terminus of the same helix as H260, increased the predicted D27 sidechain pK_a_ of D27 by 0.32 units. Proline is the most frequently observed residue type at position 256 in the PKT family MSA (Figure S6), and a similar D27 sidechain ΔpK_a_ value of +0.33 was calculated in the H256P mutant. Mutation induced increases in the predicted D27 sidechain pK_a_ at positions 256 and 260 were additive, and a ΔpK_a_ value of 0.79 was recorded for the H256Y:H260Y double mutant. Acidic aspartate and glutamate residues account for 62% of all sequence-weighted residue type counts at position 27 in the PKT family MSA (Figure S6). In contrast, in the T2A:I6T double mutant, no acidic residue sidechain pK_a_ shifts were observed in reporter positions close to the T2 (E384 and D428) and I6 (E368) mutation sites (Figure 4A).

#### (2) Substrate binding channel mutations at D56, H142, E153, D547, H548 and N549 positions

Exploratory investigation of electrostatic re-organisation arising from mutations at D547 (H_x_ = 0.37), H548 (H_x_ = 0.17) and N549 (H_x_= 0.03) positions displaying varying degrees of conservation in the mobile QN loop in the active-site principally led to the identification of H548Y as the most promising candidate. Predicted pK_a_s of the imadazole side-chains of H64, H320, and H553 histidine residues suggested in the literature to be the PKT B_2_ catalyst ^29,30,47^ were respectively shifted upwards by 0.32, 0.10 and 0.11 units in H548Y (Table 1). The ΔpK_a_ shift for the H64 sidechain able to hydrogen bond with the O1 ring oxygen atom of the bound GA-dimer (Figure 2B) is notably increased compared to the H548N mutant (Table 1). This is accompanied by a positive to negative reversal in the induced polarised electrostatic charge difference in the catalytic reaction centre, discernable from electrostatic potential difference maps for the H548N (Figure 3A) and H548Y (Figure 3B) mutants. The predicted sidechain pK_a_ of the adjacent D547 residue shifted identically by +0.21 in both mutants (Table 1). Exploitation of the differences in side-chain size and pK_a_s of aspartate (4.0) and glutamate (4.4) in model compounds ^41^ was examined as a possible means of fine tuning of the local electric field in the active-site, noting also that glutamate and aspartate are approximately equally represented at position 547 in the PKT family (Figure S6). Although the ΔpK_a_ for the residue side-chain at position 547 was raised by +0.79 in the H548Y:D547E double mutant, calculated shifts in the H64, H320, and H553 histidine side-chain pK_a_s remained almost unchanged (Table S3). Replacement of N549 with a negatively charged aspartate residue was predicted be effective in raising the sidechain pK_a_s of almost all the basic residues in the active-site, including H64 by almost one unit (Table S4). Amongst the calculated pK_a_ perturbations caused by the introduction of cysteine, serine and valine replacements at position 549, documented in Table S4, the only active-site histidine residue with a higher side-chain pK_a_ than in the H548Y mutant was H553 in the H549S mutant (ΔpK_a_ = +0.19).

The H142N catalytic centre mutant was recently isolated by Yang et al ^6^ from random pairwise residue type combination library screening at five positions (P136, H142, S440, R524 and S739) with improved catalytic efficiency towards both GA and ERU compared to the Bad.F6Pkt wild-type enzyme. Significant predicted shifts in the H64 (+0.34) and H553 (+0.21) residue side-chain pK_a_ values were observed for this mutant (Table 1), as well as an evident increase in the electric field strength close to the C2 atom of the TPP cofactor (Figure S5A). The sidechain pK_a_ of the nearby E153 residue, to which we previously drew attention ^9^, was calculated to increase by 0.21 units in an analogous fashion to that of D547 in the H548N and H548Y mutants (Table 1). Replacement of E153 by an aspartate residue in the H142N:E153D double mutant led to further calculated ΔpK_a_ increases for H64 (+0.39) and H553 (+0.27) histidine side-chains, in addition to a predicted rise of +0.11 in the H97 side-chain pK_a_, unobserved in any other mutant examined here (Table 1).

In a further attempt to influence the protein electric field in the active-site from a longer distance, D56 situated at the entrance of the substrate binding channel (Figure 2A), was replaced by an isosteric asparagine residue tolerated at this position in the PKT family (Figure S6). Although the predicted side-chain pK_a_ of the nearest acidic residue (E55) was lowered by 0.12 unit, no changes were found in the calculated side-chain pK_a_ values of histidine residues in the catalytic centre (Table S4)

### Introduction of long-range mutation increases catalytic efficiency for GA and ERU

To facilitate experimental investigation of electrostatic mutation combinations primarily aimed at promoting GA dimer ring opening, we first sought to identify an alternative background to the H548N mutant as a basis for engineering. Taking into account the size and conformational differences in preferred ring forms adopted by the GA dimer (β-furanose) and D-fructose (β-pyranose), we therefore screened the 20 active-site variants previously reported to enhance D-fructose activity^9^ for their activity on GA (see Figure S8).The PKT mutants were expressed using the *E. coli* Rosetta (DE3) plysS strain and pET-28(a)+ expression vector harbouring the corresponding gene with an N-terminal His-tag. Proteins were purified and their activity was tested using the colorimetric hydroxamate assay, that detects the phosphoketolase reaction product acetyl-P ^50,51^. The H548Y Bad.F6Pkt variant exhibited significantly improved kinetic parameters for the C_2_ aldehyde and ERU compared to both the wild-type enzyme and the H548N variant (Table 2, Supp. Inf.). The H548Y Bad.F6Pkt electrostatic mutant was therefore preferred as a more suitable single mutational variant background in addition to the wild-type enzyme for combinatorial engineering at residue positions highlighted in Figure 2A and Figure 4A.

**Table 2.** Kinetic parameters of Bad.F6Pkt wild type and selected variants. Experiments were carried out at pH 6.5, 37 °C in the presence of 50 mM sodium phosphate. If substrate saturation was not observed under the experimental conditions (n.sat.), it was not possible to determine K_M_ and consequently, the catalytic efficiency k_cat_/K_M_ could not be calculated (n.d.). Deviation of the mean is given in brackets (n ≥ 2).

| Bad.F6Pkt | D-erythrulose |  |  | glycolaldehyde |  |  |
| --- | --- | --- | --- | --- | --- | --- |
| | $V_{max}$<br>[U mg<sup>-1</sup>] | $K_M$<br>[mM] | $k_{cat}/K_M$<br>[s<sup>-1</sup> M<sup>-1</sup>] | $V_{max}$<br>[U mg<sup>-1</sup>] | $K_M$<br>[mM] | $k_{cat}/K_M$<br>[s<sup>-1</sup> M<sup>-1</sup>] |
| wild type<sup>a</sup> | 0.30<br>(± 0.09) | 36.2<br>(± 3.2) | 13.1<br>(± 4.5) | 0.41<br>(± 0.09) | n.sat. | n.d. |
| H548N<sup>a</sup> | 0.67<br>(± 0.12) | 48.7<br>(± 0.4) | 21.3<br>(± 3.6) | 0.53<br>(± 0.03) | n.sat. | n.d. |
| H548Y | 0.62<br>(± 0.08) | 26.9<br>(± 1.9) | 35.4<br>(± 4.3) | 0.41<br>(± 0.02) | 43.4<br>(± 3.5) | 14.8<br>(± 1.0) |
| H260Y:H548Y | 0.75<br>(± 0.06) | 21.2<br>(± 1.7) | 54.9<br>(± 1.9) | 0.50<br>(± 0.02) | 40.9<br>(± 1.2) | 19.0<br>(± 1.1) |
| H256Y:H260Y:H548Y | 0.59<br>(± 0.08) | 16.9<br>(± 1.4) | 49.6<br>(± 7.8) | 0.48<br>(± 0.05) | 40.3<br>(± 2.3) | 19.1<br>(± 0.8) |
| H142N<sup>b</sup> | 0.07<br>(± 0.01) | 9.0<br>(± 0.3) | 11.6<br>(± 2.0) | 0.58<br>(± 0.13) | 42.3<br>(± 3.2) | 20.6<br>(± 3.9) |
| H142N:E153D | 0.03<br>(± 0.00) | 4.9<br>(± 1.8) | 9.4<br>(± 2.4) | 0.08<br>(± 0.01) | 4.4<br>(± 0.2) | 26.3<br>(± 3.3) |

<sup>a</sup> Data published previously<sup>9</sup>. $v_{max}$ values for glycolaldehyde refer to the specific activity at a substrate concentration of 300 mM, since no saturation was observed under tested conditions.
<sup>b</sup> Data obtained in this work. Mutation H142N has been studied previously by Yang et al.<sup>6</sup>

Several of the single Bad.F6Pkt mutants computationally investigated for evidence of improved electrostatic preorganisation were then experimentally assessed for their influence on GA and ERU activity (Figure S9). Kinetic parameters determined for selected variants are shown in (Figure 5). Consistent with previous observations for F6P and fructose^9^, mutations at the highly conserved N549 position (D,C,S,V) resulted in a substantial decline in ERU and GA activity (9). While the single variants H256P, H256Y, H260Y and D56N exhibited catalytic properties comparable to those of the wild-type enzyme for both substrates, D547E increased catalytic efficiency on the C_4_ sugar by a factor of 1.9 (Figure 5) and GA activity by 33 ± 4 % (Figure S9). However, substrate saturation was not observed at concentrations of up to 300 mM GA in these single variants.

**Figure 5.**
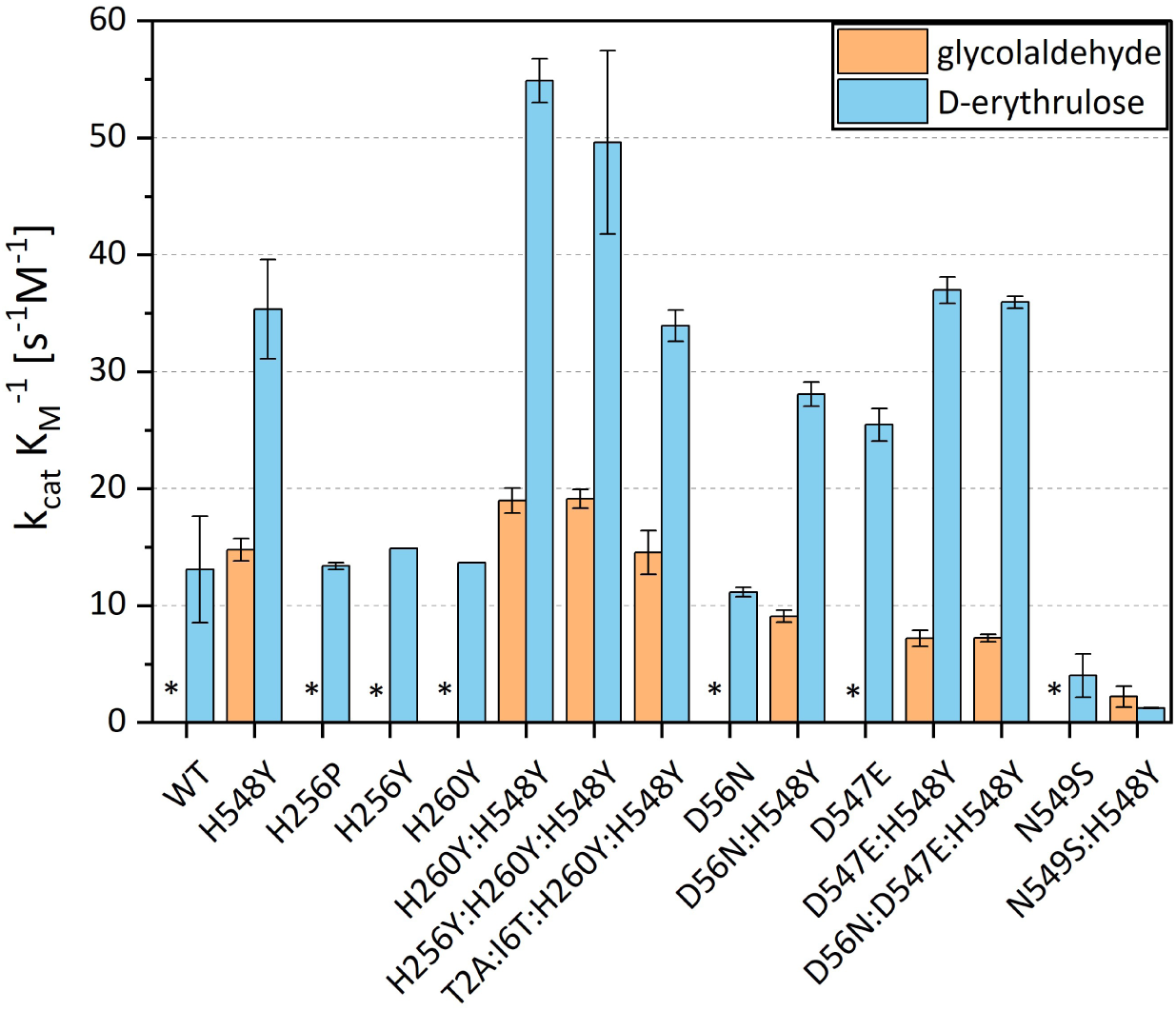
Bad.F6Pkt variant catalytic efficiency k_cat_/K_M_ for D-erythrulose and glycolaldehyde. WT – wild type. Experiments were carried out at pH 6.5, 37 °C in the presence of 50 mM sodium phosphate. Non-linear kinetic parameter fitting of experimental data to the Michaelis-Menten equation was carried out using MATLAB R2023b. In cases where substrate saturation could not be observed under experimental conditions, it was not possible to determine K_M_ or k_cat_/K_M_ (*). Error bars indicate deviation of the mean (n ≥ 2).

Substitution of H548 by a larger aromatic tyrosine residue in this study permitted the only experimental determination of a mutant k_cat_/K_M_ kinetic parameter for glycolaldehyde at this position. The catalytic efficiency of the Bad.F6Pkt variant H548Y was enhanced by 28 % for GA and 55 % for ERU through the introduction of a tyrosine residue in position H260 (Figure 5,Table 2). Comparison of the electrostatic potential difference maps for the H548Y (Figure 3B), H260Y (Figure 4B) and H548Y:H260 (Figure 3C) mutants illustrates constructive addition of the individual mutation-induced modulations of the protein electric field in the double mutant. The additional replacement of histidine 256 by a tyrosine did not influence the kinetic parameters for GA or the k_cat_/K_M_ for ERU. Nevertheless, we continued to work with the triple mutant H256Y:H260Y:H548Y, as the introduction of the third tyrosine residue significantly decreased the K_M_ value for the C_4_ sugar to 16.9 ± 1.4 mM (Table 2).

### GA affinity of Bad.F6Pkt H142N enhanced by introduction of additional E153D mutation

Yang *et al*. recently identified a Bad.F6Pkt active-site variant, H142N, which exhibited a two-fold higher V_max_ than the wild-type enzyme and a K_M_ value of 11.6 mM for GA at pH 7.5 ^6^. Although the kinetic parameters obtained under the experimental conditions of the present study differed (we applied pH 6.5), the beneficial effect of the H142N mutation on GA activity and affinity was confirmed. The catalytic efficiency of the H142N mutant for GA was found to be comparable to the H260Y:H548Y and H256Y:H260Y:H548Y variants described above (Table 2). In accordance with previous observations by Yang et al., the H142N variant exhibited a reduced K_M_ value for ERU, as well as impaired maximum activity in comparison to the wild-type enzyme (Table 2).

To explore potential synergistic effects on GA and ERU activity, we first combined the H142N mutation with previously identified substitutions shown to enhance activity or affinity towards non-phosphorylated substrates. These included H548N^9^, Q546E:N549D^9^, H548Y, H260Y:H548Y and H256Y:H260Y:H548Y. However, the majority of the resulting Bad.F6Pkt H142N variants exhibited a loss of activity towards both target substrates. The field line spatial disposition in the electrostatic potential difference map for the H142N:H548Y double mutant provides evidence of the counterproductive combined neutralisation of individual mutation-induced polarised charge in the immediate vicinity of the TPP cofactor C2 atom (Figure S5C). Only the combination of H142N with H256Y resulted in a slight increase in GA activity compared to the corresponding single variant (Figure S1).

Experimentally observed enhanced GA activity of the H142N mutant can be correlated with calculated upward shifts in the pK_a_s of putative B_2_ catalyst H64 and H553 residue sidechains (Table 1), and re-organisation of the electric field in the active-site centre (Figure S5A). In common with mutation-targeted histidine residues at positions 256, 260 and 548 that screen centres of negative charge in proximal aspartate side-chains, mutation of H142 to asparagine led to a predicted increase in the side-chain pK_a_ of the partially buried nearby E153 acidic residue (Table 1). Multiple sequence alignment weighted frequency profile analysis shows the presence of acidic glutamate residues in 93 % of PKT family sequences at this largely invariant (H_x_ = 0.14) position (Figure S6). Aspartate residues are notably poorly represented (0.8%) at position 153 in native PKTs with preferred specificities for phosphorylated substrates. E153 thus represents a promising target for further enhancement of GA activity and affinity in the Bad.F6Pkt H142N variant. Guided by a hierarchical clustered statistical correlation energy heat map for positions 142 and 153 (Figure S7) derived from MSA data (see Methods), a focused H142–E153 double mutant library was generated and assessed for GA and ERU activity. It comprised the residues D, L, P, Q, T and the wild-type in position E153, as well as asparagine, glutamate and the wild-type histidine in position 142 (Figure 6B).

**Figure 6.**
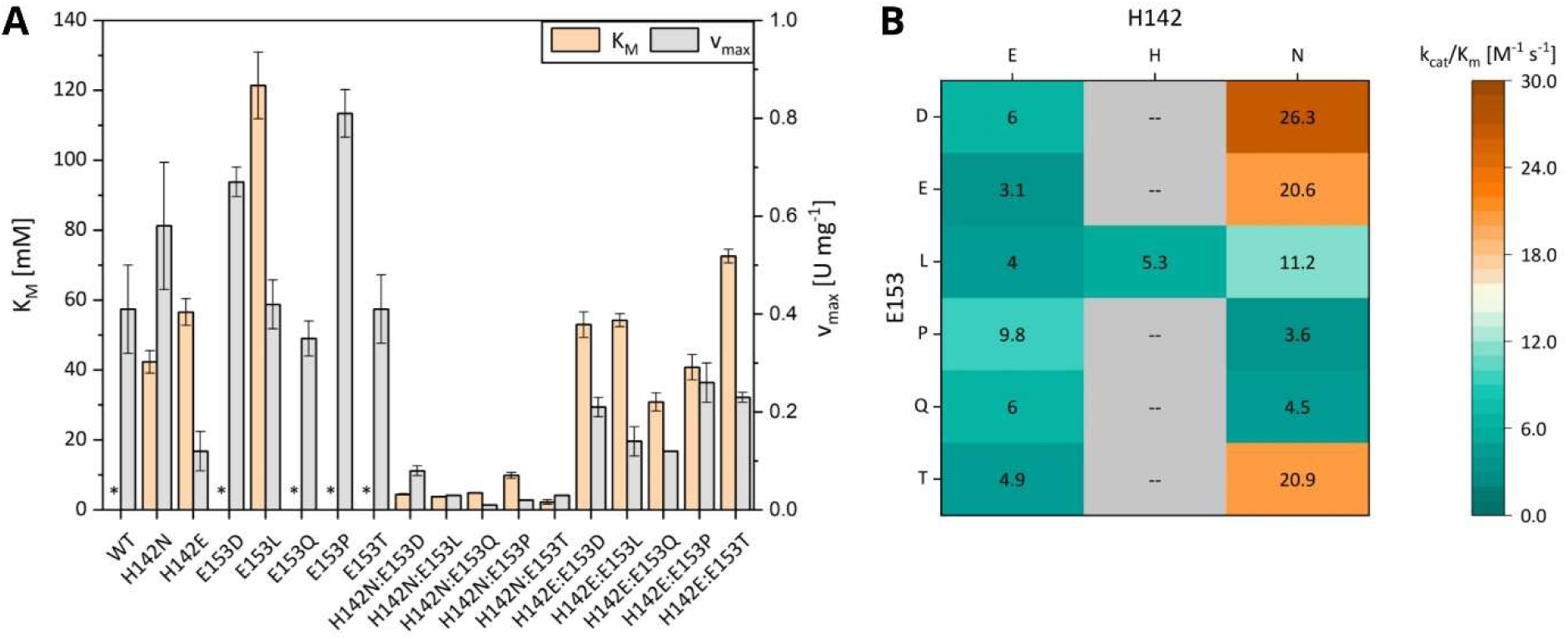
Kinetic parameters of Bad.F6Pkt H142-E153 mutants for glycolaldehyde. **A)** Michaelis-Menten constant K_M_ for glycolaldehyde and maximum specific activity V_max_. WT – wild type. Experiments were carried out at pH 6.5, 37 °C in the presence of 50 mM sodium phosphate. Non-linear kinetic parameter fitting of experimental data to the Michaelis-Menten equation was performed using MATLAB R2023b. If no substrate saturation was observed under tested conditions, K_M_ could not be determined (*) and the specific activity at a substrate concentration of 300 mM is given as V_max_. Error bars indicate deviation of the mean (n ≥ 2). **B)** Catalytic efficiency k_cat_/K_M_ for glycolaldehyde. – In the abscence of a K_M_ estimate, k_cat_/K_M_ cannot be determined. Values represent the mean of at least two replicates.

As observed for the Bad.F6Pkt H142N single mutant (Table 2), all tested H142-E153 double mutants exhibited a decrease in ERU activity of at least 60 % in comparison to the wild-type enzyme, along with a significant reduction in catalytic efficiency towards the C_4_ sugar (data not shown). The introduction of either glutamate or asparagine in position H142 of the Bad.F6Pkt E153 variants was found to significantly decrease V_max_ for GA (Figure 6A). However, the asparagine residue conferred a pronounced improvement in GA affinity, with K_M_ values for the H142N:E153 variants approximately one order of magnitude lower than those of the corresponding single mutants (Figure 6A). Among these, only Bad.F6Pkt H142N:E153D exhibited higher catalytic efficiency for the C_2_ aldehyde than the H142N single mutant, which was increased by approximately 28% (Figure 6B, Table 2). A K_M_ value of 4.4 ± 0.2 mM was observed for this double mutant, corresponding to a 10-fold higher GA affinity compared to H142N (Figure 6A, Table 2). A marked reduction in V_max_ accompanying the significant lowering of the Michaelis constant is characteristic of the preferential stabilisation of the substrate ground-state over the reaction transition-state. Comparison of the electrostatic potential difference maps shown in Figure S5 for the H142N (A) and H142N:E153D (B) mutations reveals the clear displacement of high-density field line zones associated with improved catalytic activity away from the enzyme reaction centre in the double mutant.

### Bad.F6Pkt variants improve the performance of ATP synthesis from either GA or ERU

Practical advantages provided by the newly constructed enzymes were investigated using in vitro ATP regeneration from GA or ERU as a model system. The reaction system produced *sn*-glycerol 3-phosphate (G3P) from glycerol and ATP using glycerol kinase from *Cellulomonas sp.* NT3060 (Cs.GlpK). Acetate kinase from *Geobacillus stearothermophilus* (Gs.AckA) was employed to enable ATP regeneration from ADP. Both enzymes were previously found to be highly active and stable under synthesis-like conditions^9,52,53^. PKT activity in the reaction systems was modulated by employing either the Bad.F6Pkt wild-type enzyme or selected variants with enhanced kinetic parameters for ERU and/or GA at a concentration of 2 mg mL^-1^. The ATP-dependent G3P production was assessed using an initial concentration of 25 mM ERU or GA, 1 mM ADP and an excess of glycerol and phosphate (50 mM, respectively).

First, we evaluated the G3P synthesis from GA by the GA-specific Bad.F6Pkt variants H256Y:H260Y:H548Y, H142N and H142N:E153D in comparison with the wild-type enzyme. The reaction employing the Bad.F6Pkt H142N:E153D variant proceeded for 10 hours, resulting in a final G3P concentration of 19.5 ± 0.9 mM and a slightly increased yield (0.8 ± 0.08 mol_G3P_ mol_GA_^-1^) compared to the wild-type (0.7 ± 0.00 mol_G3P_ mol_GA_^-1^)(Figure 7). Further, it was observed that reactions containing the wild-type PKT or the H256Y:H260Y:H548Y mutant ceased G3P production after six hours, reaching final product concentrations of 17.9 ± 0.5 mM and 22.7 ± 0.3 mM, respectively (Figure 7B). This incomplete substrate conversion may be attributed to PKT inactivation in the presence of GA^25^. In contrast, complete conversion of the limiting substrate GA to G3P was achieved within four hours using the Bad.F6Pkt H142N variant (Figure 7), thus demonstrating the feasibility of *in vitro* ATP regeneration from the non-phosphorylated C_2_ aldehyde. Notably, the utilization of Bad.F6Pkt H142N also resulted in the highest maximum G3P productivity of 17.7 ± 0.1 mM h^-1^, consistent with its comparatively high V_max_ (Figure 7A,Table 2). Despite its improved GA affinity, the double Bad.F6Pkt mutant H142N:E153D possessed a significantly lower kinetic turnover number compared to H142N (Table 2), reflected by its inability to outperform the single variant in terms of ATP regeneration from an initial GA concentration of 25 mM.

**Figure 7.**
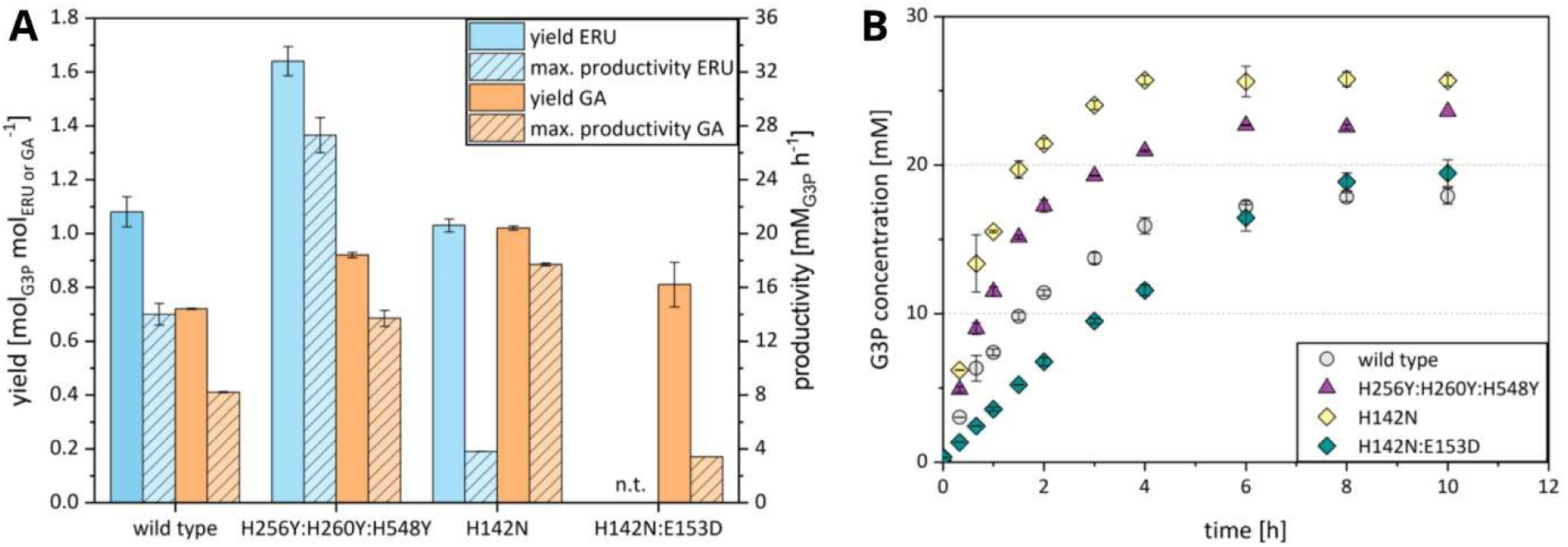
***In vitro* ATP regeneration from either ERU (blue) or GA (orange) by a phosphoketolase. A)** *In vitro sn*-glycerol 3-phosphate (G3P) yield and productivity with ATP regenerated from ERU or GA using different Bad.F6Pkt variants or the wild-type enzyme. **B)** G3P production by different Bad.F6Pkt variants or the wild-type enzyme using GA as substrate. A) + B) Experiments were carried out at pH 7.0, 37 °C in the presence of 50 mM sodium phosphate and glycerol, 0.8 mM TPP, 4 mM MgCl_2_, 1 mM ADP and a starting concentration of 25 mM D-erythrulose or glycolaldehyde. Phosphoketolase activity was modulated by employing either wild-type Bad.F6Pkt or indicated variants at a concentration of 2 mg mL^-1^. n.t. – not tested. Data represent mean and deviation of biological duplicates.

After demonstrating the GA-based ATP regeneration system, we sought to investigate whether the *in vitro* synthesis of up to two mol ATP from one mole ERU via the intermediate GA (as shown in Figure 1) is feasible. Here, the performance of the ERU-specific Bad.F6Pkt variant H256Y:H260Y:H548Y was compared with the wild-type enzyme. The single mutant H142N was included as a control, as it enabled complete conversion of GA that is present as an intermediate in the ERU-based ATP regeneration system. ATP regeneration from 25 mM ERU using the Bad.F6Pkt H256Y:H260Y:H548Y variant enabled the synthesis of up to 40.2 ± 1.3 mM G3P within six hours (Figure S11A). This corresponds to a yield of 1.64 ± 0.05 mol_G3P_ mol_ERU_^-1^, demonstrating that both the PKT-catalysed conversion of ERU to acetyl-P and the subsequent degradation of the resulting GA intermediate contributed to ATP regeneration in this system (see Figure 1). Furthermore, a maximum productivity of 27.3 ± 1.3 mM h^-1^ was achieved with the Bad.F6Pkt triple mutant, which is nearly twice that observed with the wild-type enzyme (Figure 7A). In contrast, utilization of the wild-type Bad.F6Pkt or the H142N variant resulted in significantly lower productivity and smaller yields of approximately 1 mol_G3P_ mol ^-1^, as G3P synthesis ceased after six and eight hours, respectively (Figure 7A, Figure S11A). It is noteworthy that only the use of the H256Y:H260Y:H548Y mutant identified in this study led to near-complete consumption of the C_4_ sugar substrate (residual ERU concentration was 2.1 ± 0.1 mM). In the same reaction, up to 7 mM GA were accumulated (Figure S11B).

### Coupling of threose aldolase and threose isomerase with ERU-specific PKT enables rapid conversion of GA and increases the ATP yield

Following the successful demonstration of ATP regeneration from ERU and GA, respectively, we set out to investigate whether incorporating a D-threose aldolase and D-threose isomerase into the reaction system could enhance ATP yield from GA by accelerating the degradation of the enzyme-inactivating C_2_ aldehyde (see Figure 1). The *E. coli* fructose 6-phosphate aldolase (Ec.FsaA) variant L107Y:A219G was employed for the conversion of two molecules of GA to one molecule of D-threose, due to its reported high activity and affinity for GA^26^. The D-threose isomerase activity was provided by L-rhamnose isomerase from *Pseudomonas stutzeri* (Ps.LrhI)^54^, for which kinetic parameters were determined (Table 3). *In vitro* G3P synthesis from 15 mM GA by glycerol kinase, acetate kinase and the Bad.F6Pkt variant H256Y:H260Y:H548Y was evaluated in the presence and absence of threose aldolase and isomerase. The PKT triple mutant was utilized in both reactions at a concentration of 0.2 mg mL^-1^, due to its beneficial properties regarding ERU and GA, respectively.

**Table 3.** Properties of enzymes used for *in vitro* acetyl-P synthesis. Unless otherwise stated, kinetic parameter data obtained in this work represent mean values and standard deviation from at least two experiments carried out at 37°C and pH 7.0. The equilibrium constant K_eq_ was determined using the web tool eQuilibrator 3.0 ^55^ with the following settings: pH 7.0, pMg 3.0 and ionic strength 0.25 M. n.sat. – no substrate saturation.

| Enzyme | Go.Gox0313 | Ms.Nox | Ec.FsaA L107Y:A129G | Ps.Lrhl |
| --- | --- | --- | --- | --- |
| Organism | *Gluconobacter oxydans* | *Methanobrevibacter smithii* | *Escherichia coli* | *Pseudomonas stutzeri* |
| Function | ethylene glycol dehydrogenase | NADH oxidase | D-threose aldolase | threose isomerase |
| $V_{max}$ [U mg<sup>-1</sup>] | 0.14 ± 0.02<sup>a</sup> | 0.03 ± 0.00 | 5.3 ± 0.3<sup>d</sup> | 1.0 ± 0.2 |
| $K_M$ [mM] | n.sat. (EG) | 0.05 – 0.06 (NADH)<sup>c</sup> | 0.09 ± 0.01 (GA, donor)<sup>d</sup> | 25.7 ± 3.2 (D-threose) |
|  | 0.06 (NAD<sup>+</sup>)<sup>b</sup> | 0.02 (O<sub>2</sub>)<sup>c</sup> | 2.1 ± 0.3<sup>d</sup> (GA, acceptor) |  |
| $K_{eq}$ | $7.0 \times 10^{-5}$ | $1.1 \times 10^{40}$ | 670 | 3 |

<sup>a</sup> $V_{max}$ refers to the specific activity at an ethylene glycol concentration of 100 mM, since no saturation was observed under tested conditions
<sup>b</sup> parameters estimated at pH 8.5 and 30 °C<sup>56</sup>
<sup>c</sup> parameters estimated at pH 7.2<sup>57</sup>
<sup>d</sup> parameters estimated at pH 7.5 and 25 °C<sup>26,58</sup>

Without conversion of GA to threose and ERU, G3P synthesis ceased after 8 hours (Figure 8A). With approximately half of the initial GA concentration remaining, the observed loss of PKT activity can be attributed to enzyme inactivation caused by the aldehyde (^25^, Table S6). In contrast, in the presence of threose aldolase and isomerase, complete consumption of GA occurred within less than 30 minutes. The transient accumulation of the intermediates threose and ERU (Figure 8B) indicated that the coupling of threose aldolase and isomerase enabled the conversion of GA to ERU through the intended pathway (Figure 1). The performance of the reaction system was significantly improved, as witnessed by the G3P yield which increased from 0.47 ± 0.06 mol_G3P_ mol_GA_^-1^ to 0.69 ± 0.06 mol_G3P_ mol_GA_^-1^ (Figure 8). However, a complete conversion of GA to G3P was not achieved and the reaction ceased with 0.8 ± 0.0 mM D-threose and 3.6 ± 0.3 mM ERU remaining in the reaction mix. As previously reported for its diastereomer D-erythrose^9^, D-threose also irreversibly inactivates the PKT enzyme, reducing its half-life from 144 h^9^ to 19.7 h at a concentration of 10 mM. Although the pseudo-first order rate constant for inactivation by D-threose is approximately one order of magnitude lower than that of GA when present at the same molar concentration (Table S6), the incomplete conversion of GA to G3P can likely be attributed to irreversible inactivation of the PKT by D-threose.

**Figure 8.**
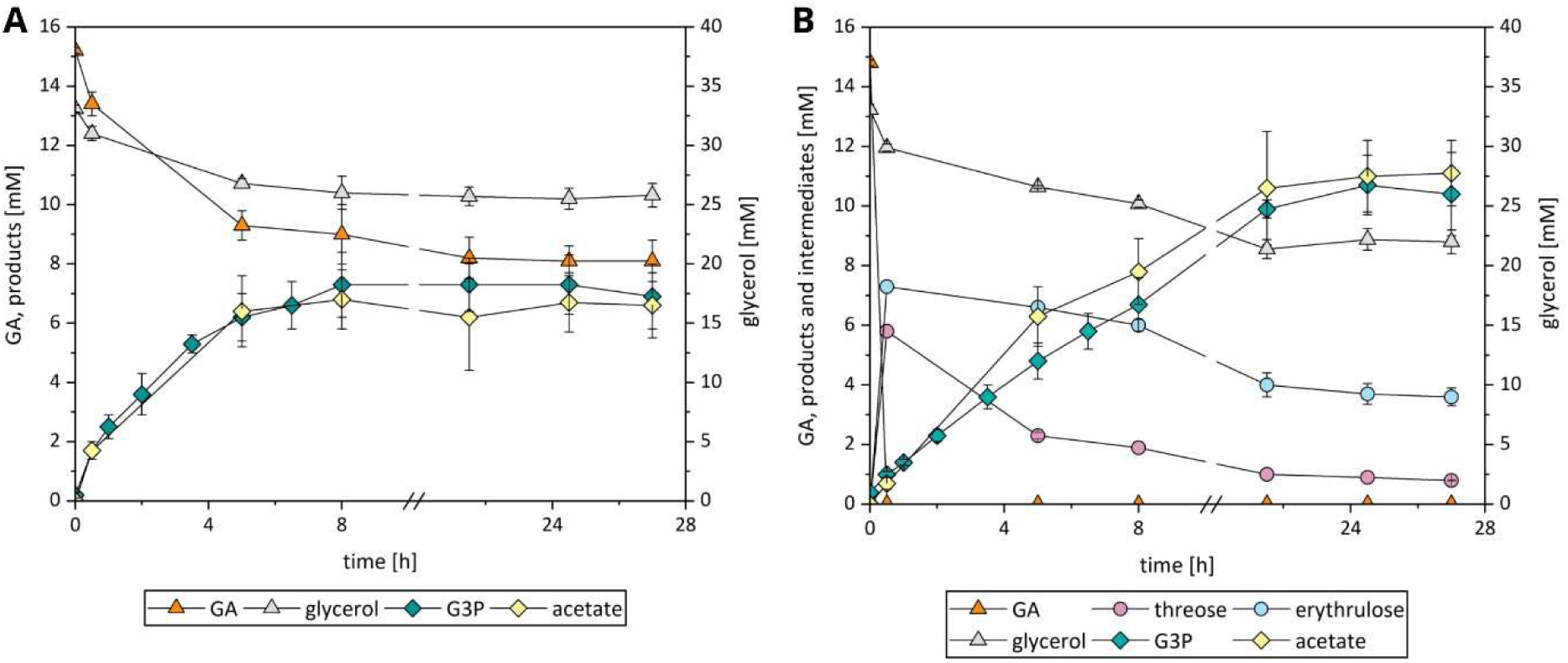
Glycolaldehyde (GA) conversion in phosphoketolase-based reaction systems for in vitro ATP regeneration. **A)** Course of substrate and product concentrations over time using 0.2 mg mL^-1^ of Bad.F6Pkt H256Y:H260Y:H548Y for acetyl-P production. **B)** Course of substrate, intermediate and product concentrations over time using 0.2 mg mL^-1^ of Bad.F6Pkt H256Y:H260Y:H548Y in combination with 0.33 mg mL^-1^ Ec.FsaA L107Y:A129G and 2.5 mg mL^-1^ Ps.LrhI for *acetyl-P*production. A) + B) Experiments were carried out at pH 7.0, 37 °C in the presence of 32 mM sodium phosphate and glycerol, 0.8 mM TPP, 4 mM MgCl_2_, 1 mM ADP and a starting concentration of 15 mM glycolaldehyde. Data represent mean and deviation of biological duplicates.

### ATP regeneration from ethylene glycol is enhanced by improved PKT variants

As illustrated in Figure 1, GA is proposed to be generated *in situ* from the less toxic^10^ and less expensive bulk chemical ethylene glycol (EG) in the envisioned ATP regeneration system(s). Based on literature reports, EG dehydrogenase from *Gluconobacter oxidans* (Go.Gox0313)^56,59,60^ and NADH oxidase from *Methanobrevibacter smithii* (Ms.Nox)^57^ were selected for the oxidation of EG to GA and characterized in terms of their activity at synthesis-like conditions (pH 7.0, 37 °C)(Table 3). As both enzymes exhibited adequate activities, they were subsequently combined with a GA-specific PKT variant, acetate kinase and glycerol kinase to assess the functionality of the *in vitro* reaction system for ATP regeneration from EG (as shown in Figure 1). The two Bad.F6Pkt variants with the highest catalytic efficiency for GA (H142N and H142N:E153D) were compared to the wild-type enzyme, in order to investigate whether employing a GA-specific PKT could enhance the reaction system performance. Given that GA reduction is thermodynamically favoured over EG oxidation (Table 3, ^10,56^), EG was supplied in excess (100 mM), while glycerol (22 mM) served as the limiting substrate for *in vitro* G3P synthesis.

Compared to the wild-type Bad.F6Pkt, employing the GA-specific variants H142N or H142N:E153D resulted in a two-fold higher G3P productivity during the initial seven hours of the reaction (Figure 9), reaching a maximum of 0.78 mM h^-1^. The utilization of the Bad.F6Pkt variant H142N increased the overall G3P yield from 0.28 ± 0.02 mol_G3P_ mol ^-1^ (wild-type enzyme) to 0.52 ± 0.02 mol_G3P_ mol ^-1^. In contrast, a precipitate was observed in reaction mixtures containing Bad.F6Pkt H142N:E153D after 6 hours, which, together with halted conversion of GA (inferred from stagnating acetate and G3P concentrations), indicates reduced stability of the double mutant under synthesis conditions. The highest GA accumulation occurred in the reaction employing the wild type PKT (1.1 ± 0.1 mM), followed by the double and the single mutant (0.8 ± 0.2 mM and 0.4 ± 0.1 mM).

**Figure 9.**
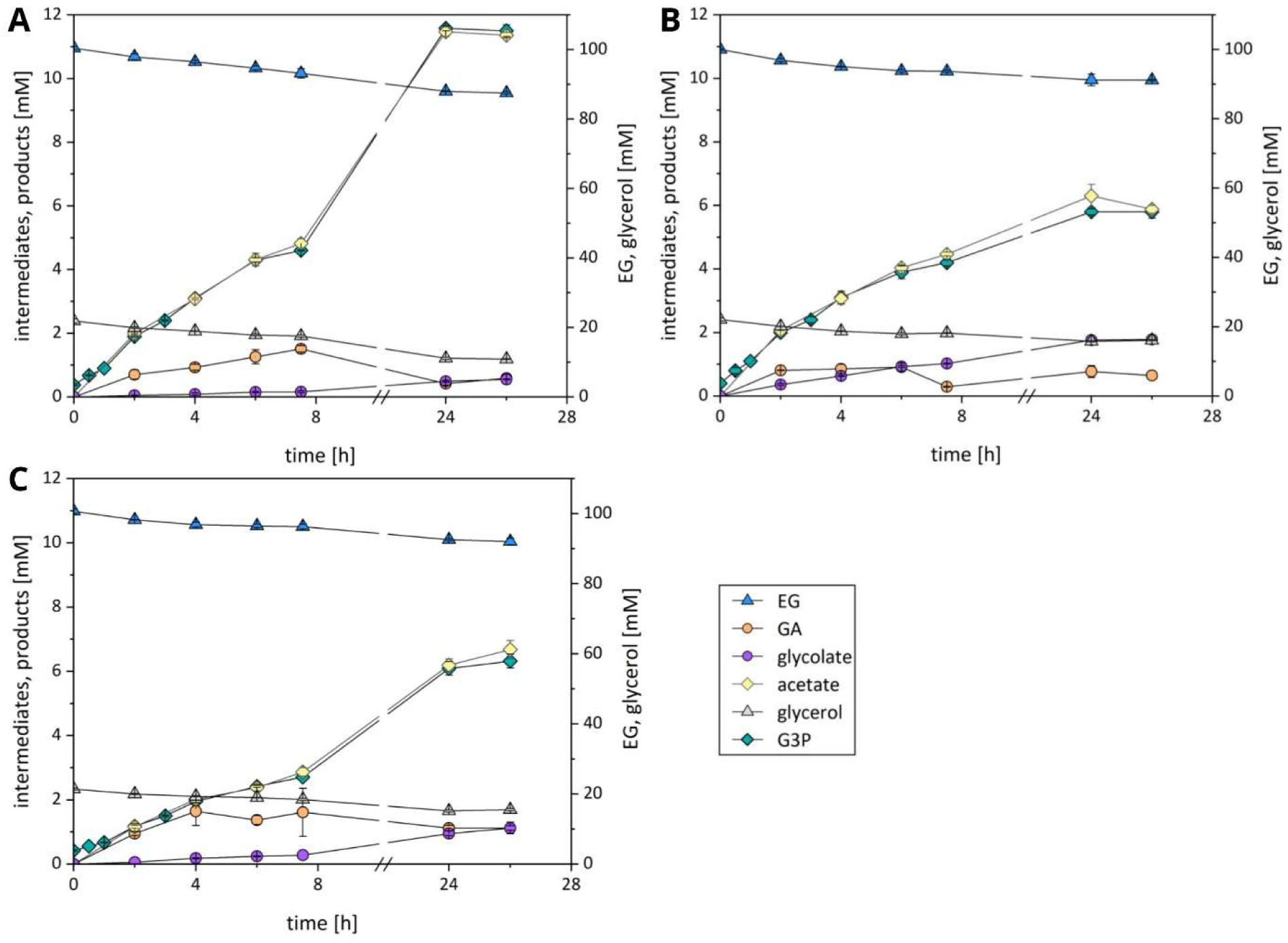
***In vitro* sn-glycerol 3-phosphate (G3P) synthesis with a reaction system for ATP regeneration based on ethylene glycol.** Substrate consumption, accumulation of intermediates and (by-) products during *in vitro* G3P synthesis using 2 mg mL^-1^ of **A)** Bad.F6Pkt H142N, **B)** Bad.F6Pkt H142N:E153D or **C)** the wild-type phosphoketolase enzyme in combination with 0.5 mg mL^-1^ Go.Gox0313 and 1.4 mg mL^-1^ Ms.Nox for acetyl-P production. Experiments were carried out at pH 7.0, 37 °C and 200 rpm in the presence of 0.8 mM TPP, 4 mM MgCl_2_, 2 mM NAD^+^ and 1 mM ADP, with initial concentrations of 100 mM ethylene glycol, 32 mM sodium phosphate and 22 mM glycerol. Data represent mean and deviation of biological duplicates.

## Discussion

In this study, we successfully applied a rational electrostatic enzyme engineering design approach to improve the affinity and catalytic efficiency of PKT from *B. adolescentis* for the non-phosphorylated substrates ERU and GA. Additionally, we implemented an *in vitro* ATP regeneration system, which is based on the oxidation of the bulk chemical EG to GA and subsequent production of the high-energy phosphoryl donor acetyl-P by an engineered PKT. Efficient conversion of the cell-toxic intermediate GA was achieved *in vitro* either by employing a GA-specific PKT variant or, alternatively, by coupling D-threose aldolase and D-threose isomerase with an ERU-specific PKT in the presence of inorganic phosphate.

Engineering of the wild-type PKT was focussed on promotion of *in situ* ring-opening of the cyclic GA dimer and the generation of the reactive monomeric form ^7^ to enhance substrate specificity for the C_2_ aldehyde. Molecular modelling of the Bad.F6Pkt–TPP complex with bound cyclic GA dimer, combined with multiple sequence alignment analysis and calculations of side-chain pK_a_ shifts at ionizable protein residues, enabled the identification of electrostatic mutations in two distinct regions of the enzyme that could facilitate GA dimer ring opening. Whereas the wild-type Bad.F6Pkt showed no substrate saturation up to 300 mM GA, substitution of the active-site residue H548 by a larger aromatic tyrosine permitted experimental determination of a k_cat_/K_M_ kinetic parameter for GA and increased the catalytic efficiency for ERU by a factor of 2.7. Improved catalytic performance towards GA and ERU could be correlated with predicted upward shifts in H64, H320 and H553 side-chain pK_a_ values that appear to at least partially mirror those induced in native PKTs by the terminal phosphate groups of cognate X5P and F6P phosphorylated substrates. Introduction of the long-range mutation H260Y into the H548Y background further enhanced catalytic efficiency for both GA and ERU, as predicted by the constructive modulation of the protein electric field in the enzyme catalytic centre (Figure 3). With a 4.2-fold higher k_cat_/K_M_ than the wild-type enzyme, this double mutant exhibited the highest catalytic efficiency for the open-chain ERU reported to date^6,9^. This may be tentatively attributed to a lowering of TS energy barriers for deprotonation of the bound TPP-ERU covalent adduct or the dehydration step by the double mutant. Additional replacement of histidine 256 by a tyrosine did not affect GA kinetics but slightly improved ERU binding without significantly altering k_cat_/K_M_.

Both the affinity and catalytic efficiency towards GA of the PKT variant H142N identified by Yang et al.^6^ were improved by substituting glutamate at position 153 with aspartate in a double mutant. However, although H142N:E153D displayed a suitably low K_M_ of 4 mM for biological application purposes, its maximum activity towards GA decreased by approximately seven-fold. Much of the binding energy gained in stabilising the substrate ground-state would not appear to be available for reaction TS stabilisation. We are currently seeking to further stabilise TS binding in this mutant through the engineering of additional differentially orientated induced dipoles at distances further removed from the catalytic centre.

The feasibility of cell-free ATP regeneration from GA with a stoichiometry of 1 mol ATP per mol of C_2_ aldehyde was demonstrated using the PKT single mutant H142N together with acetate kinase and glycerol kinase for the production of G3P. The PKT double mutant H142N:E153D did not outperform the single variant in terms of ATP regeneration from an initial GA concentration of 25 mM, likely due to its significantly lower V_max_. These results indicate that the single variant is better suited for cell-free reaction systems, whereas the double mutant with its ten-fold lower K_M_ for GA may be advantageous for *in vivo* applications (e.g. acetyl-CoA synthesis from EG^6–8^), where only very low concentrations of the toxic C_2_ aldehyde are tolerated^10^.

The *in vitro* synthesis of up to 1.64 mol ATP per mol ERU via the intermediate GA was shown using the ERU-specific Bad.F6Pkt variant H256Y:H260Y:H548Y which outperformed the wild type enzyme and the H142N variant. This experiment laid the grounds for a strategy that aimed at rapidly removing the enzyme-inactivating GA from the reaction buffer by converting it to D-threose and eventually to the PKT substrate D-erythrulose through a reaction sequence catalysed by D-threose aldolase and D-threose isomerase. We demonstrated that the use of a D-threose aldolase and D-threose isomerase in combination with the ERU-specific PKT triple mutant H256Y:H260Y:H548Y indeed enabled rapid GA consumption and an associated increase in product yield compared to direct utilisation of GA by the PKT. However, despite these positive effects incomplete substrate conversion was observed, likely due to accumulation of the aldose intermediate D-threose, which exhibited an inactivating effect on the PKT enzyme. Nevertheless, this reaction system may hold potential for *in vivo* detoxification of GA during production of the platform chemical acetyl-CoA from EG or methanol ^8,10^.

Cell-free ATP regeneration from EG was achieved by combining an EG dehydrogenase and NADH oxidase with a GA-specific PKT variant, acetate kinase and glycerol kinase. Employing the Bad.F6Pkt variant H142N or H142N:E153D resulted in a two-fold higher maximum productivity compared to the wild-type enzyme, emphasizing the importance of a GA-specific PKT for efficient EG-based ATP-regeneration. Due to reduced stability of the double mutant under synthesis conditions, the highest yield (0.52 ± 0.02 mol_G3P_ mol ^-1^) was obtained with the single mutant H142N.

In conclusion, this study demonstrates that catalytic field engineering is an effective strategy to expand PKT substrate diversity towards smaller, non-phosphorylated substrates. The resultant variants provide for improved biocatalysis in GA-dependent metabolic pathways and cell-free ATP regeneration systems based on the low-cost feedstock EG.

## Methods

### Reagents and chemicals

Unless otherwise stated, all chemicals were purchased from Merck KGaA (Darmstadt, Germany). Commercial kits for plasmid DNA isolation, gel DNA extraction and PCR clean-up were obtained from New England Biolabs (Frankfurt am Main, Germany) and used according to the manufacturer’s instructions. Synthetic genes were purchased from BioCat GmbH (Heidelberg, Germany) or Thermo Fisher Scientific Inc (Waltham, MA, USA), while genomic DNAs were obtained from the German Collection of Microorganisms and Cell Cultures GmbH (DSMZ, Braunschweig, Germany). Genewiz Germany GmbH (Leipzig, Germany) or Eurofins Genomics (Ebersberg, Germany) carried out DNA sanger sequencing.

### Plasmid construction and site-directed mutagenesis of *Bad.f6Pkt*

Plasmids used in this study to express the N-terminally 6x-His-tagged target genes are listed in Table S5. Mutations were introduced individually to the *Bad.f6pkt* gene by single site-directed mutagenesis using PCR-based method described by Zheng and colleagues^61^. The PCR reaction was performed with Phusion™ High-Fidelity DNA polymerase (NEB), 62.5 ng of template vector and primers listed in Supporting Information 2. After digestion of residual template DNA by DpnI restriction endonuclease (NEB), plasmids were transformed into chemically competent *E. coli* NEB 5-alpha cells (NEB). DNA sanger sequencing was used to verify respective mutations.

### Protein production

Expression of N-terminally 6x-His-tagged enzymes was carried out in *E. coli* strains harboring the corresponding pET-28a(+) expression vector. Gs.AckA and Ec.FsaA L107Y:A129G were produced in *E. coli* BL21 (DE3) cells (New England Biolabs), whereas *E. coli* Rosetta(DE3) plysS (Merck KGaA) served as the host strain for all other enzymes, including Go.Gox0313 and Ms.Nox. The expression procedure was based on the method described previously ^8,9^. Briefly, cultures were grown in LB medium at 37 °C to mid-log phase, induced with 1 mM isopropyl-β-D-thiogalactopyranoside (IPTG) and incubated at 25 °C for 20 h. Production of Bad.F6Pkt and Ms.Nox was carried out using the auto-inducing medium ZYM-5052^62^, which was inoculated with an initial optical density at 600 nm of 0.05, prior to incubation at 25 °C for 24 h.

### Protein purification, quantification and storage

Protein purification was performed following the procedure described previously^9^. Briefly, cell-free crude extract was applied to 0.35 mL Talon™ Cobalt affinity resin (Cytiva, Marlborough, USA) and incubated at room temperature and 20 rpm rotation (RevolverTM, Labnet, Edison, NJ, USA) for 1 h. The resin was washed twice with buffer (10 mM KH_2_PO_4_, 300 mM NaCl, pH 7.5) before bound His-tagged enzymes were eluted using 0.5 mL elution buffer (10 mM KH_2_PO_4_, 300 mM NaCl, 500 mM imidazole, pH 7.0). Buffer exchange of the eluate was performed with an Amicon Ultra-0.5 centrifugal filter unit (pore size 10 kDa; Merck Millipore, USA). By centrifugation (1000 g, 4 °C, 2 min.) the protein fraction was recovered in 0.4 mL storage buffer (10 mM KH_2_PO_4_, 300 mM NaCl, pH 7.0) and stored at 4 °C until further use. Purified Ms.Nox solution was supplemented with 1 mM flavin adenine dinucleotide (FAD) prior to storage. Protein concentrations were determined by the Bradford assay (Roti®-Quant, Carl Roth, Karlsruhe, Germany) using bovine serum albumin (0–100 µg mL⁻¹) as the calibration standard.

### Multiple sequence alignment residue and residue-pair amino acid frequency analyses

Position-dependent residue type frequencies and relative Shannon information entropy measures of residue variability (H_X_) in an HMM multiple sequence alignment (MSA) comprising 1019 PKT enzyme family sequences ^9^ were calculated using SEQUESTER ^63^ from sequence-weighted data counts to reduce phylogenetic bias. MSA sequence weights were calculated according to the method of Morcos *et al*. ^64^ for *q* =20 native amino acid residue types with an identity cut-off of 80%, before normalisation to sum to the total number of sequences. H_X_ values at position X in the Bad.F6Pkt sequence fall within the limits of zero, corresponding to a fully conserved residue position, and one, for a position showing no intrinsic residue type preference.

Sequence weighted frequency counting was also applied in the calculation of statistical correlation energy measures, − ln 𝑔(𝑋_i_ , 𝑌_j_), of the pairwise dependence of residue type (*i* =1,20) at position X on the residue type (*j*=1,20) at position Y in the MSA. The correlation 𝑔(𝑋*_i_* , 𝑌*_j_*) is given by 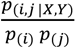, where 𝑝_(i,j |X,Y)_ is the conditional pairwise probability of observing (*i*,*j*) residue types respectively at positions X and Y. Background probabilities of residue type occurrence (𝑝_(i)_, 𝑝_(j)_) were taken as mean frequency values for all proteins ^65^. Data frequency seeding was implemented to address dual concerns of data sparseness and rare or forbidden events.Seed counts were set as 𝑞^2^ 𝑝_(i)_ 𝑝_(j)_, corresponding to an average of a single phantom count per residue type combination. Favourable or most frequently observed residue type combinations at a given pair of residue positions are associated with negative values of − ln 𝑔(𝑋_i_ , 𝑌_j_), whereas positive values of the statistical energy measure correspond to unfavourable or infrequently observed combinations. A zero score implies the lack of dependence between the two residue types.

### Molecular modelling of Bad.F6Pkt-TPP complex with glycoaldehyde cyclic dimer

The close structural resemblance of the five-membered 2-hydroxymethyl-4-hydroxy-1,3-dioxalane (Figure S1) and the β-furanose ring form of D-fructose was exploited to construct a model of the glycolaldehyde cyclic dimer bound to the Bad.F6Pkt-TPP complex. Docking of a HF/6-31G* geometry-optimised 2-hydroxymethyl-4-hydroxy-1,3-dioxalane ligand molecule was carried out by superposition of the C2, O1, C5, C4 and O3 atom centres on the structurally equivalent C2, C3, C4, C5 and O5 β-furanose ring skeleton atoms in the previously described modelled complex of D-fructose with Bad.F6Pkt and Mg-TPP ^9^. Following excision of the D-fructose ligand co-ordinates, the resultant structure was then energy minimised in the presence of non-overlapping X-ray crystallographic water molecules abstracted from the *Bifidobacterium breve* Pkt H320A mutant homologue template (PDB code 3ahi) used in the modelling of Bad.F6Pkt ^9^. The *ff99SB* Amber molecular mechanics force field ^66^ was used for protein atoms, and the electrostatic model comprised a distance-dependent dielectric constant with ε=4. GAFF force field parameters ^67^ and partial charges for the 4’-aminopyrimidine tautomeric form of TPP^2-68^ were as previously reported ^9^. A complete list of GAFF atom types and RESP ^69^ partial charges for 2-hydroxymethyl-4-hydroxy-1,3-dioxalane is provided in Table S1 of the Supplementary Information. All Hartree-Fock (HF) quantum chemical calculations were performed using GAMESS ^70^.

### Protein Continuum Electrostatics Calculations

Ionisable (Asp, Glu, His, Lys and Arg) protein residue pK_a_ calculations were performed on wild type Bad.F6Pkt and minimally disturbed mutant enzyme model co-ordinate sets using DelPhiPKa version 2.3 ^42,43^. Parameters for the reference dielectric constant (ε) in the Gaussian distribution formula for the protein interior and the variance (σ) of the Gaussian distribution were left at their respective default values of 8.0 and 0.70. A 12 Å cluster threshold delimitation was used. The C++ software code for automated hydrogen atom placement was modified before compilation of the program to permit adjustment of protein mainchain secondary amide N-H bond lengths from 1.5 Å to 1.0 Å. The default AMBER force-field was used in all calculations. Reference side-chain pK_a_ values for L-glutamic acid (4.00) and L-histidine (6.50) in water were respectively altered in the steering data input file to 4.31 ^71^ and 6.20 ^45^ in closer agreement with experimental values. Simulated residue titrations in the pH range 0 to 14 were carried out at pH intervals of 0.5.

Side-chain mutations were initially introduced into the dimeric wild type Bad.F6Pkt enzyme model using the COOT interactive molecular graphics and modelling package ^72^ before the application of a single round of energy minimisation. Any significant conformational changes to the protein residue side-chain geometry arising from energy minimisation were re-introduced into the starting atomic co-ordinate set. A single Mg-TPP cofactor microstate was included in all Poisson-Boltzmann continuum electrostatic calculations. Atomic charges for the TPP^2-^ 4’-aminopyrimidine tautomer were taken from Lie *et al*. ^68^ and the corresponding radii from the standard Bondi compendium ^73^ with minor adjustments.

Poisson-Boltzmann electrostatic potential (φ(**r**)) grid calculations on PQR-formatted co-ordinate sets generated by DelPhiPKa comprising prevalent protein residue ionisation states at pH 7, were performed using APBS version 3.4.1 ^74^ with an internal dielectric constant (ε) of 4 and at an ionic strength of 0.15M. DX-formatted Δφ(**r**) electrostatic potential difference volumetric maps, calculated by subtraction of mutant and wild-type enzyme electrostatic potential (φ(**r**)) grids using the *dxmath* APBS conversion utility, were displayed as isopotential surfaces and field lines in the VMD graphics program ^75^.

### Enzymatic assays

Infinite M200 PRO plate reader (Tecan AG, Switzerland), I-control software (version 3.8.2.0, Tecan AG) and 96-well flat bottomed microplates (Thermo Fisher Scientific Inc) were used to determine the activities of purified enzymes. One unit of enzyme activity (U) is defined as synthesis of 1 micro mole of product per minute. Experimental data were fitted to the Michaelis–Menten or substrate inhibition model using the Curve Fitting Tool in MATLAB (version R2023b, MathWorks Inc., Natick, MA, USA) to determine the enzyme’s Michaelis constant (K_M_) and maximum reaction velocity (v_max_).

Enzymatic activity of phosphoketolase was determined as described previously^9^ using the hydroxamate assay^50,51^ to detect the product acetyl-P. The enzymatic reaction was carried out at 37 °C and pH 6.5 in the presence of 50 mM KH_2_PO_4_, 3.8 mM MgCl_2_ and 0.8 mM TPP using either 200 mM GA or 25 mM ERU as the substrate. For the estimation of kinetic parameters, specific activity was measured at substrate concentrations ranging from 0 – 300 mM (GA) or 0 – 100 mM (ERU).

D-threose isomerase activity was quantified by the detection of ERU using D-erythrulose reductase from *Gallus gallus* (Gg.DER) in a coupled assay adapted from the procedure described by Morii *et. al*^76^. The reductase enzyme was expressed, purified and stored as described previously ^9^. The reaction mixture with at total volume of 250 µL contained 100 mM HEPES buffer pH 7.0, 1 mM MnCl_2_, 0.2 mM NADH (dissolved in 10 mM NaOH) and 8 µg mL^-1^ Gg.DER and appropriate quantities of the purified isomerase enzyme. The enzymatic reaction was initiated by addition of D-threose in concentrations ranging from 0 – 80 mM and monitored continuously at 37 °C by the detection of NADH at 340 nm.

The activity of ethylene glycol dehydrogenase in the oxidative direction was determined by monitoring the reduction of NAD^+^ at 340 nm and 37 °C. The reaction mixture contained 100 mM HEPES buffer pH 7.0, 0.5 mM NAD^+^ and appropriate amounts of purified enzyme. Reactions were started by the addition of ethylene glycol at variable concentrations (0 – 100 mM).

NADH oxidase activity was quantified by monitoring the consumption of NADH at 340 nm and 37 °C. Appropriate quantities of the purified enzyme were present in 100 mM HEPES buffer pH 7.0 and the reaction was initiated by the addition of 0.25 mM NADH.

### *In vitro sn*-glycerol 3-phosphate (G3P) synthesis

The reaction mixture for *in vitro* G3P synthesis contained 200 mM HEPES buffer (pH 7.0), 1 mM ATP, 0.8 mM TPP, 4 mM MgCl_2_, 32 mM sodium phosphate (pH 7.0) and glycerol, 5 µg mL^-1^ Gs.AckA and 0.3 mg mL^-1^ Cs.GlpK, unless otherwise stated. The following enzymes were added as required: Bad.F6Pkt (variants), Go.Gox0313, Ms.Nox, Ec.FsaA L107Y:A129G and Ps.LrhI. After a preliminary incubation at 37 °C for 5 minutes, the reactions was initiated by the addition of either 25 mM ERU, 15 or 25 mM GA or 100 mM EG. Each reaction (1 mL total volume) was performed in a 2 mL tube and incubated at 37 °C for 26 h. Reactions containing EG as the substrate were supplemented with 2 mM NAD^+^ and shaken at 220 rpm. Samples of 50 µL were obtained at regular intervals and subsequently diluted 1:4 in 0.1 M HCl to quench enzymatic activity, followed by immediate cooling on ice. For real-time, offline monitoring of G3P production using the coupled enzymatic assay described below, a portion of each quenched sample was further diluted in 0.1 M HEPES buffer (pH 7.5). The remaining aliquots were stored at −20 °C until further analysis.

### Quantification of substrates, intermediates and products

The target product G3P was quantified using the photometric method described previously^9^. Briefly, G3P is converted by *sn*-glycerol-3-phosphate oxidase (G3Pox, Creative Enzymes, USA) to dihydroxyacetone phosphate and hydrogen peroxide. In the presence of 4-aminoantipyrine (4-AAP) and sodium 3,5-dichloro-2-hydroxybenzenesulfonate (DHBS), the subsequent peroxidase-catalyzed reaction consumes one mole of H₂O₂ to yield one mole of water and quinone imine, which is detected photometrically at 520 nm. Absorbance was monitored at 30 °C until signal stabilization and G3P concentration was determined from the absorbance difference between the sample and a control reaction lacking G3Pox.

Ethylene glycol, glycolaldehyde, C4 sugars, glycolate, glycerol and acetate were quantified by high-performance liquid chromatography (UltiMate3000, Thermo Fisher Scientific, USA) equipped with a DAD-detector (DAD-3000(RS), Thermo Fisher Scientific, USA), a RI-detector (RefractoMax 520, ERC, Germany) and an autosampler with a temperature set to 6°C. After appropriate dilution in ddH_2_O, samples were filtered (0.2 μm PTFE filter) before injection of 20 µL to the Rezex^TM^ ROA-Organic Acid H^+^ (8 %) column (Phenomenex). Analytes were eluted with 0.5 mM H_2_SO_4_ at a flow rate of 0.5 mL min^–1^ and a temperature of 65 °C.

## Author contributions

CMT performed the electrostatics calculations, and carried out the computational structure and sequence variation analyses. CMT, FK and TW developed the enzyme engineering strategy. FK and KR performed molecular biology work and carried out enzymatic assays. FK performed *in vitro* product syntheses. Data curation and visualization were done by FK. FK, JOK, DO, AL and TW designed experimental setups. FK, TW and CMT wrote the paper, which was reviewed by JOK, DO and AL. TW conceived the cascade and supervised the project.

## Supporting Information

**Supporting Information 1** – Protein electrostatics calculations, MSA analysis, plasmids used and additional results

**Supporting Information 2** – Primers used in this study

## Acknowledgments

We would like to thank our colleagues Cláudio J.R. Frazão, Lisa Rothe and Sebastian Schulz for providing the plasmids pET28a_Ms. nox, pET28a_Go.gox0313 and pET28a_Ec.fsaA_L107Y:A129G.

