## Supplementary Information 1 for "Electrostatic Engineering of Phosphoketolase Enhances Activity on Small Non-phosphorylated Sugars and Improves Cell-Free ATP Regeneration from Inexpensive C_2_-Substrates"

### Phosphoketolase reaction mechanism on GA and ERU

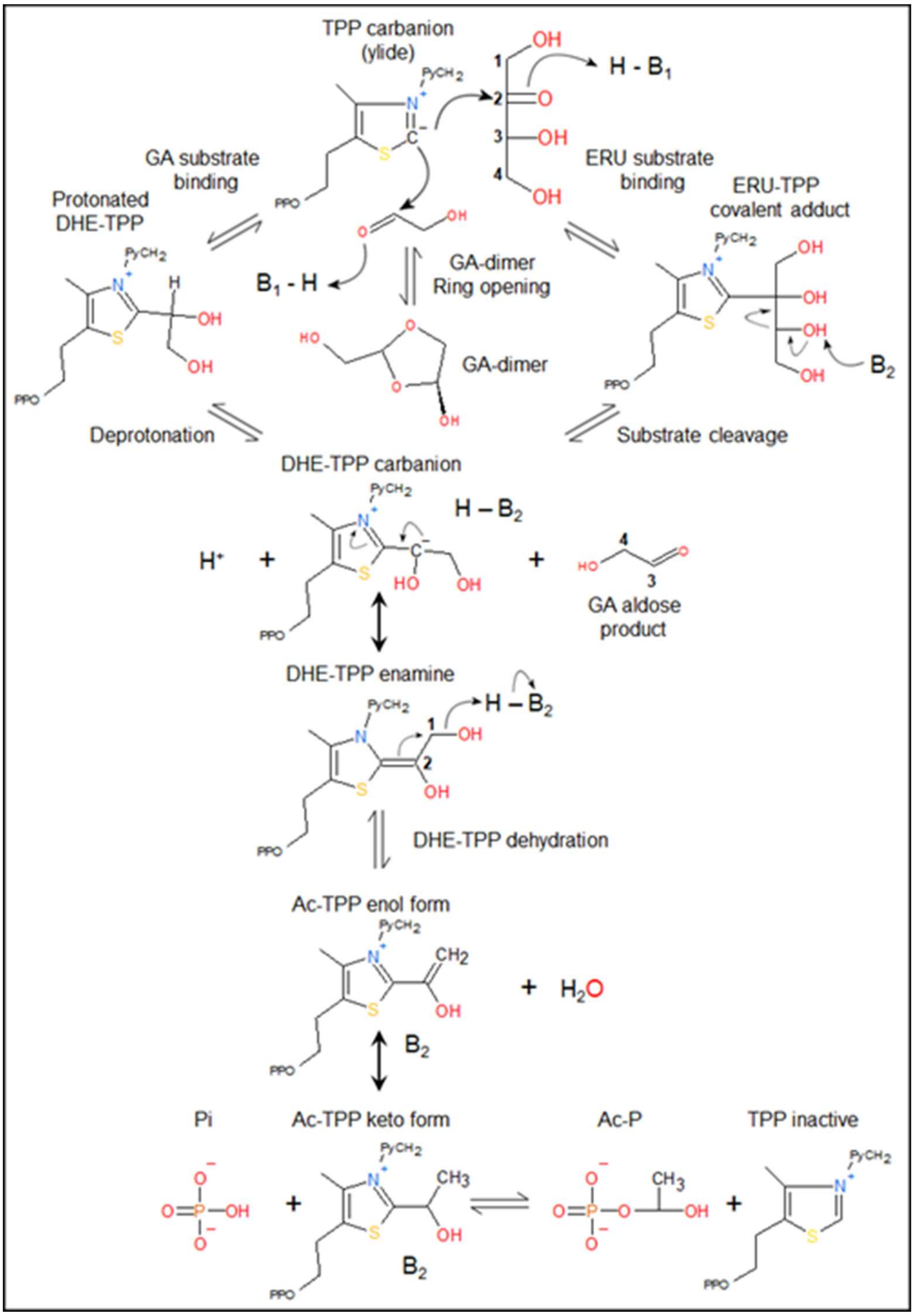


**Scheme S1 Phosphoketolase reaction mechanisms on glycoaldehyde and D-erythrulose.** Key reaction (resonance) intermediates and catalytic steps implicated in alternative non-phosphorylated substrate-dependent mechanisms[1–7] are indicated. Abbreviations: GA, glycoaldehyde; GA-dimer, 2-hydroxymethyl-4-hydroxy-1,3-dioxolane; ERU, D-erythrulose; TPP, thiamine pyrophosphate; Py, aminopyrimidine; PPO, diphosphate; DHE-, dihydroxyethyl; Ac-, acetyl; Ac-P, acetyl phosphate; Pi, phosphate. Attributed PKT catalytic base residues in the literature: $B_1$, H553; $B_2$, H64, H320, H553 (Bad.F6Pkt sequence numbering). Participation of H64 in enzyme-catalysed ring opening of the cyclic GA dimer for *in situ* generation of the monomeric GA species can be inferred from hydrogen bonding interactions with the GA dimer in the modelled complex of Bad.F6Pkt and the Mg-TPP cofactor (Figure 2B).

### Glycoaldehyde cyclic dimer structure and force field parameters

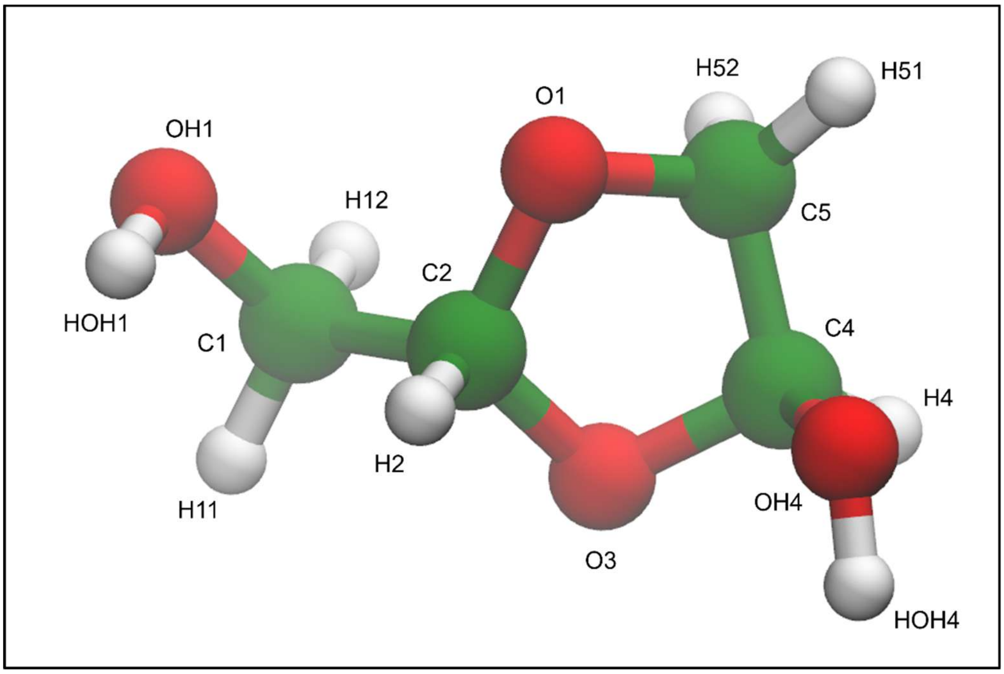


**Figure S1 2-Hydroxymethyl-4-hydroxy-1,3-dioxolane glycoaldehyde cyclic dimer structure.**

**Table S1 Partial charges and GAFF atom types for 2-hydroxymethyl-4-hydroxy-1,3-dioxolane.** RESP partial charges[8] were obtained from fitting to the HF/6-31G* quantum chemical electrostatic potential of the fully geometry-optimised structure. Atom names are shown in Supporting Information Figure S1.

| Atom Name | Partial Charge | GAFF Atom Type |
|---|---|---|
| C1 | 0.0575 | c3 |
| C2 | 0.3147 | c3 |
| C4 | 0.4906 | c3 |
| C5 | 0.0662 | c3 |
| O1 | -0.3952 | os |
| OH1 | -0.6468 | oh |
| O3 | -0.5095 | os |
| OH4 | -0.7181 | oh |
| H11 | 0.0968 | h1 |
| H12 | 0.0968 | h1 |
| HOH1 | 0.4219 | ho |
| H2 | 0.0899 | h2 |
| H4 | 0.0255 | h2 |
| HOH4 | 0.4579 | ho |
| H51 | 0.0759 | h1 |
| H52 | 0.0759 | h1 |

### Protein electrostatics calculations

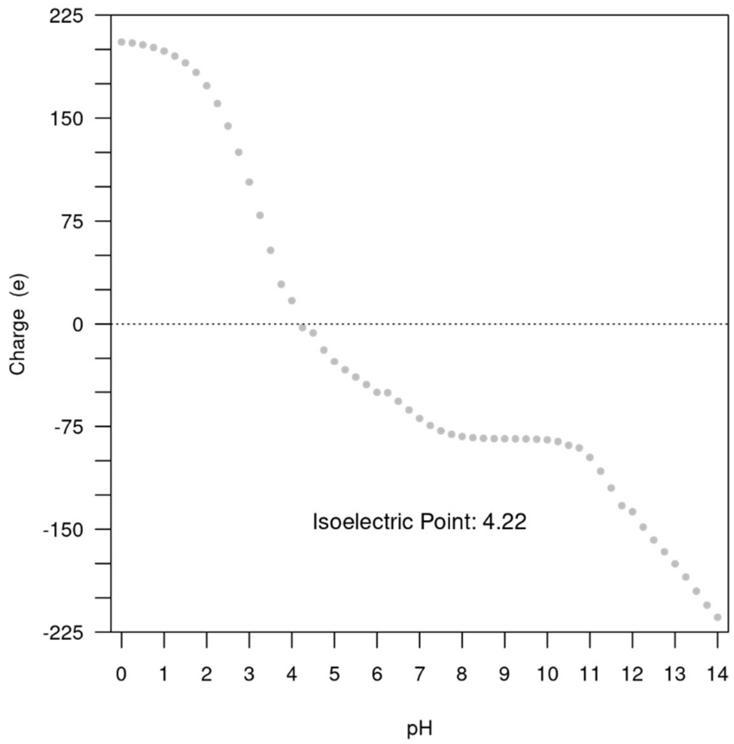


**Figure S2 Simulated titration curve for wild-type Bad.F6Pkt.** Ionisable residue p$K_a$s were calculated using DelPhiPKa[9,10] at pH intervals of 0.25 at an ionic strength of 0.15 mM as described in Methods. Each monomeric subunit carries a net charge of -36 at pH 7.

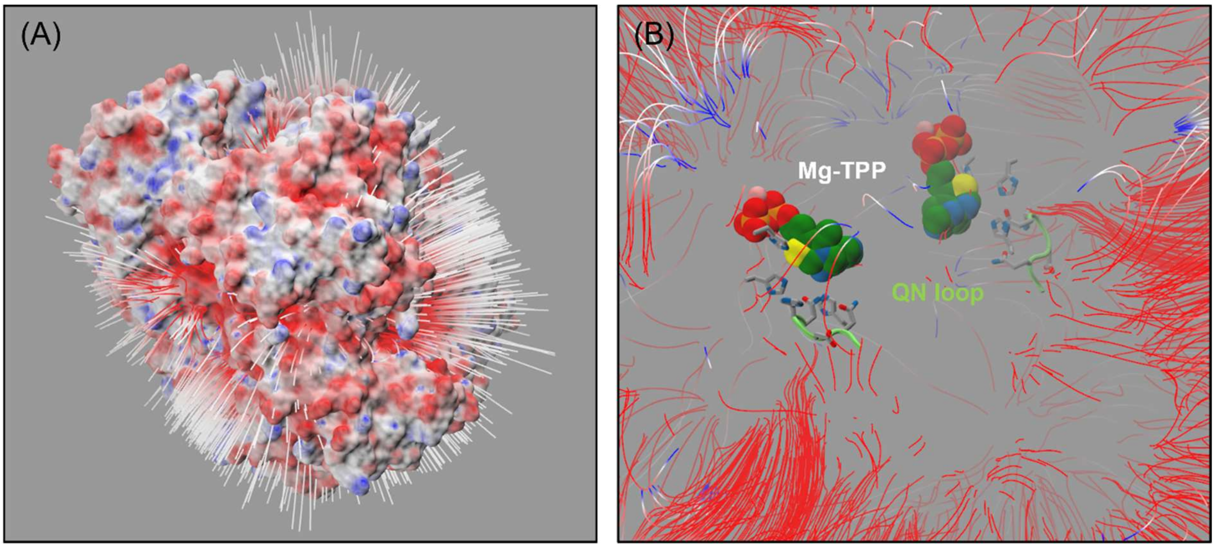


**Figure S3 Electrostatic field in wild-type Bad.F6Pkt.** Field lines are coloured according to electrostatic potential, with starting points in blue (positive potential) and end points in red (negative potential). **(A) View from protein exterior.** Electrostatic potential mapped onto the solvent-excluded surface is colour scaled at ±10 kT/e thresholds. **(B) Protein interior.** TPP cofactors and histidine (64, 320 and 553) residue side-chains in the catalytic centres of the dimeric enzyme molecule in the same orientation are respectively shown as van der Waals sphere and stick-form representations. The mainchain backbones of the QN (Q546, D547, H548, N549) active-site loops are highlighted as lime green coloured cartoon tubes, and the loop side-chains depicted as sticks with grey coloured carbon atoms.

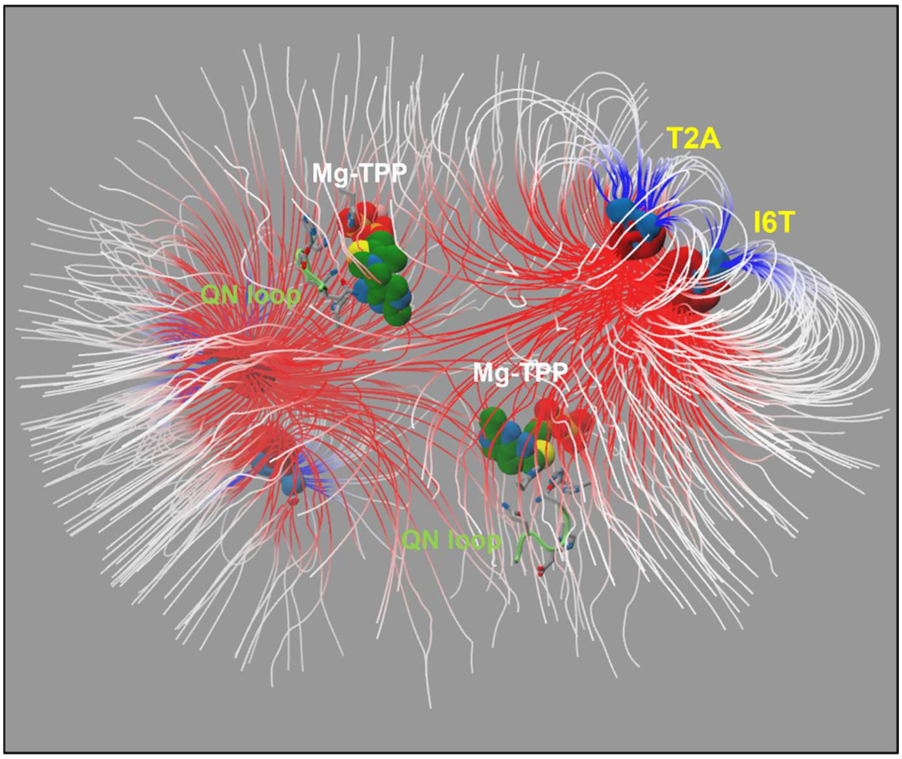


**Figure S4 Electrostatic potential difference grid map for Bad.F6Pkt T2A:I6T double mutant.** Difference electrostatic potential (Δϕ(**r**)), calculated relative to wild-type Bad.F6Pkt, is displayed as solid rendered (±2 kT/e) contoured isosurfaces, respectively coloured in blue or red for positive or negative Δϕ(**r**). Electrostatic field lines are coloured according to the electrostatic difference potential. The two TPP cofactor molecules and their bound $Mg^{2+}$ ions in the enzyme dimer are shown as van der Waals spheres, coloured according to element type: carbon, green; nitrogen, blue; oxygen, red; sulphur, yellow; phosphorus, orange; magnesium, pink. Active-site histidine (64, 320 and 553) and (Q546, D547, H548, N549) QN loop residue side-chains are shown as in stick-form with grey-coloured carbons. The QN loop mainchains are depicted in cartoon form as lime green coloured tubes.

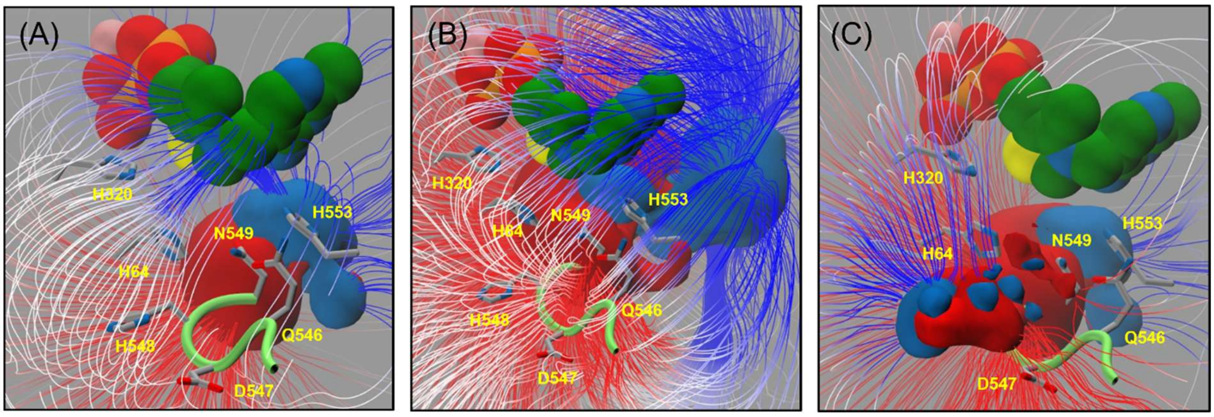


**Figure S5 Electrostatic potential difference grid maps for Bad.F6Pkt for H142N mutation carrying variants. (A)** H142N **(B)** H142N:E153D. **(C)** H142N:H548Y. Difference electrostatic potential (Δϕ(**r**)), calculated relative to wild-type Bad.F6Pkt, is shown as solid rendered ±4 kT/e contoured isosurfaces, coloured in blue or red for positive or negative Δϕ(**r**), respectively. Electrostatic field lines directed away from positive charge difference sources are coloured according to the electrostatic difference potential due to mutation. The Mg-TPP cofactor is shown as van der Waals spheres with green coloured carbon atoms. Active-site histidine (64, 320 and 553) and QN loop (Q546, D547, H548, N549) residue side-chains are shown as in stick-form with grey-coloured carbons. The QN loop mainchain is depicted in cartoon form as a lime green tube.

**Table S2. Computed residue p$K_a$ values in Bad.F6Pkt N-terminal and Asp26 region mutants.** Calculations at ionisable residue positions were carried on modelled wild-type and mutant Bad.F6Pkt complexes with Mg-TPP using DelPhiPKa. Tabulated predicted p$K_a$ values at selected positions are averages over both chains in the dimeric enzyme. For clarity, only induced p$K_a$ shifts relative to the wild type enzyme ($\Delta pK_a$) of ≥ ±0.1 in magnitude are indicated below in parentheses. RSA, side-chain residue solvent accessible surface area relative to that in corresponding extended conformation Ala-XXX-Ala model reference tri-peptide structure [11]. NA, not applicable.

| Bad.F6Pkt Residue Position | RSA (%) | Wild Type | H256P | H256Y | H260Y | H256Y: H260Y | T2A: I6T | T2A: I6T: H260Y |
|---|---|---|---|---|---|---|---|---|
| D26 | 13 | 1.97 | 2.30 (+0.33) | 2.29 (+0.32) | 2.42 (+0.45) | 2.76 (+0.79) | 1.97 | 2.50 (+0.53) |
| H59 | 26 | 5.55 | 5.55 | 5.55 | 5.55 | 5.55 | 5.55 | 5.55 |
| H64 | 2 | 5.08 | 5.09 | 5.09 | 5.09 | 5.09 | 5.08 | 5.04 |
| H97 | 0 | 6.09 | 6.09 | 6.09 | 6.10 | 6.10 | 6.09 | 6.05 |
| H142 | 2 | 4.90 | 4.91 | 4.91 | 4.91 | 4.91 | 4.90 | 4.84 |
| E153 | 21 | 2.23 | 2.23 | 2.23 | 2.23 | 2.23 | 2.23 | 2.22 |
| H256 | 46 | 6.08 | NA | NA | 6.22 (+0.14) | NA | 6.08 | 6.20 (+0.12) |
| H260 | 0 | 6.38 | 6.55 (+0.17) | 6.54 (+0.16) | NA | NA | 6.38 | NA |
| K300 | 0 | 11.86 | 11.86 | 11.86 | 11.86 | 11.78 | 11.86 | 11.88 |
| H320 | 2 | 4.64 | 4.65 | 4.65 | 4.65 | 4.66 | 4.64 | 4.71 |
| E368 | 55 | 4.00 | 4.00 | 4.00 | 4.00 | 4.00 | 3.98 | 3.97 |
| E384 | 67 | 3.98 | 3.98 | 3.98 | 3.98 | 3.98 | 3.97 | 3.99 |
| D428 | 47 | 3.43 | 3.43 | 3.43 | 3.43 | 3.43 | 3.45 | 3.41 |
| E437 | 1 | 2.23 | 2.24 | 2.24 | 2.24 | 2.24 | 2.24 | 2.24 |
| R442 | 31 | 13.16 | 13.16 | 13.13 | 13.17 | 13.16 | 13.16 | 13.17 |
| E479 | 1 | 2.58 | 2.58 | 2.50 | 2.58 | 2.58 | 2.58 | 2.58 |
| D547 | 26 | 1.19 | 1.19 | 1.19 | 1.19 | 1.19 | 1.18 | 1.18 |
| H548 | 29 | 5.52 | 5.52 | 5.52 | 5.52 | 5.52 | 5.52 | 5.53 |
| H553 | 0 | 4.93 | 4.93 | 4.93 | 4.93 | 4.93 | 4.93 | 4.96 |
| K605 | 19 | 10.76 | 10.76 | 10.71 | 10.76 | 10.76 | 10.76 | 10.79 |

**Table S3 Computed residue p$K_a$ values in Bad.F6Pkt QN loop and catalytic-centre mutants**. Calculations at ionisable residue positions were carried on modelled wild-type and mutant Bad.F6Pkt complexes with Mg-TPP using DelPhiPKa. Tabulated predicted p$K_a$ values at selected positions are averages over both chains in the dimeric enzyme. For clarity, only Induced p$K_a$ shifts relative to the wild type enzyme (Δp$K_a$) of ≥ ±0.1 in magnitude are indicated below in parentheses. RSA, side-chain residue solvent accessible surface area relative to that in corresponding extended conformation Ala-XXX-Ala model reference tri-peptide structure [11]. NA, not applicable.

| Bad.F6Pkt Residue Position | RSA (%) | Wild Type | E153D | H142N:E153Q | H142N:H548Y | D547E | D547E:H548Y |
|---|---|---|---|---|---|---|---|
| D26 | 13 | 1.97 | 1.97 | 1.97 | 1.98 | 1.97 | 1.97 |
| E55 | 80 | 3.94 | 3.94 | 3.94 | 3.95 | 3.94 | 3.94 |
| D56 | 14 | 3.09 | 3.09 | 3.12 | 3.13 | 3.09 | 3.12 |
| H59 | 26 | 5.55 | 5.55 | 5.56 | 5.64 | 5.58 | 5.64 |
| H64 | 2 | 5.08 | 5.09 | 5.29 (+0.21) | 5.69 (+0.61) | 5.07 | 5.39 (+0.31) |
| H97 | 0 | 6.09 | 6.12 | 6.07 | 6.12 | 6.09 | 6.09 |
| H142 | 2 | 4.90 | 5.18 (+0.28) | NA | NA | 4.89 | 4.94 |
| E153 | 21 | 2.23 | 1.55 (-0.68) | NA | 2.45 (+0.22) | 2.23 | 2.25 |
| H256 | 46 | 6.08 | 6.08 | 6.08 | 6.08 | 6.08 | 6.08 |
| H260 | 0 | 6.38 | 6.38 | 6.38 | 6.42 | 6.38 | 6.38 |
| K300 | 0 | 11.86 | 11.87 | 11.82 | 11.85 | 11.86 | 11.86 |
| H320 | 2 | 4.64 | 4.64 | 4.66 | 4.78 (+0.14) | 4.63 | 4.74 (+0.10) |
| E437 | 1 | 2.23 | 2.24 | 2.30 | 2.37 (+0.14) | 2.24 | 2.29 |
| R442 | 31 | 13.16 | 13.16 | 13.15 | 13.15 | 13.21 | 13.17 |
| E479 | 1 | 2.58 | 2.58 | 2.63 | 2.64 | 2.58 | 2.59 |
| D547 | 26 | 1.19 | 1.19 | 1.22 | 1.47 (+0.28) | 1.98 (+0.79) | 2.25 (+1.06) |
| H548 | 29 | 5.52 | 5.51 | 5.52 | NA | 5.55 | NA |
| H553 | 0 | 4.93 | 4.89 | 4.92 | 5.26 (+0.34) | 4.87 | 4.97 |
| K605 | 19 | 10.76 | 10.76 | 10.73 | 10.80 | 10.77 | 10.80 |

**Table S4 Computed residue p$K_a$ values in Bad.F6Pkt substrate binding channel mutants at positions D56 and N549.** Calculations at ionisable residue positions were carried on modelled wild-type and mutant Bad.F6Pkt complexes with Mg-TPP using DelPhiPKa. Tabulated predicted p$K_a$ values at selected positions are averages over both chains in the dimeric enzyme. For clarity, only Induced p$K_a$ shifts relative to the wild type enzyme (Δp$K_a$) of ≥ ±0.1 in magnitude are indicated below in parentheses. RSA, side-chain residue solvent accessible surface area relative to that in corresponding extended conformation Ala-XXX-Ala model reference tri-peptide structure [11]. NA, not applicable.

| Bad.F6Pkt Residue Position | RSA (%) | Wild Type | D56N | N549D | N549C | N549S | N549V |
|---|---|---|---|---|---|---|---|
| D26 | 13 | 1.97 | 1.97 | 1.97 | 1.97 | 1.97 | 1.97 |
| E55 | 80 | 3.94 | 3.82 (-0.12) | 3.94 | 3.94 | 3.94 | 3.94 |
| D56 | 14 | 3.09 | NA | 3.09 | 3.09 | 3.10 | 3.09 |
| H59 | 26 | 5.55 | 5.51 | 5.58 | 5.55 | 5.55 | 5.55 |
| H64 | 2 | 5.08 | 5.08 | 6.04 (+0.96) | 5.15 | 4.93 (-0.15) | 4.93 (-0.15) |
| H97 | 0 | 6.09 | 6.09 | 6.10 | 6.06 | 6.07 | 6.06 |
| H142 | 2 | 4.90 | 4.90 | 5.21 (+0.31) | 4.71 (-0.19) | 4.74 (-0.16) | 4.65 (-0.25) |
| E153 | 21 | 2.23 | 2.23 | 2.30 | 2.22 | 2.23 | 2.23 |
| H256 | 46 | 6.08 | 6.07 | 6.08 | 6.08 | 6.08 | 6.08 |
| H260 | 0 | 6.38 | 6.38 | 6.38 | 6.38 | 6.38 | 6.38 |
| K300 | 0 | 11.86 | 11.86 | 11.95 | 11.85 | 11.85 | 11.84 |
| H320 | 2 | 4.64 | 4.64 | 4.77 (+0.13) | 4.64 | 4.65 | 4.64 |
| E437 | 1 | 2.23 | 2.24 | 2.42 (+0.18) | 2.25 | 2.25 | 2.25 |
| R442 | 31 | 13.16 | 13.16 | 13.30 (+0.14) | 13.17 | 13.16 | 13.16 |
| E479 | 1 | 2.58 | 2.58 | 2.58 | 2.58 | 2.58 | 2.57 |
| D547 | 26 | 1.19 | 1.19 | 1.27 | 1.20 | 1.20 | 1.20 |
| H548 | 29 | 5.52 | 5.53 | 5.67 (+0.15) | 5.55 | 5.52 | 5.54 |
| N549 | 5 | NA | NA | 0.36 | NA | NA | NA |
| H553 | 0 | 4.93 | 4.93 | 6.10 (+1.17) | 4.76 (-0.17) | 5.12 (+0.19) | 4.87 |
| K605 | 19 | 10.76 | 10.76 | 11.04 (+0.28) | 10.81 | 10.80 | 10.80 |

### Multiple sequence alignment analysis

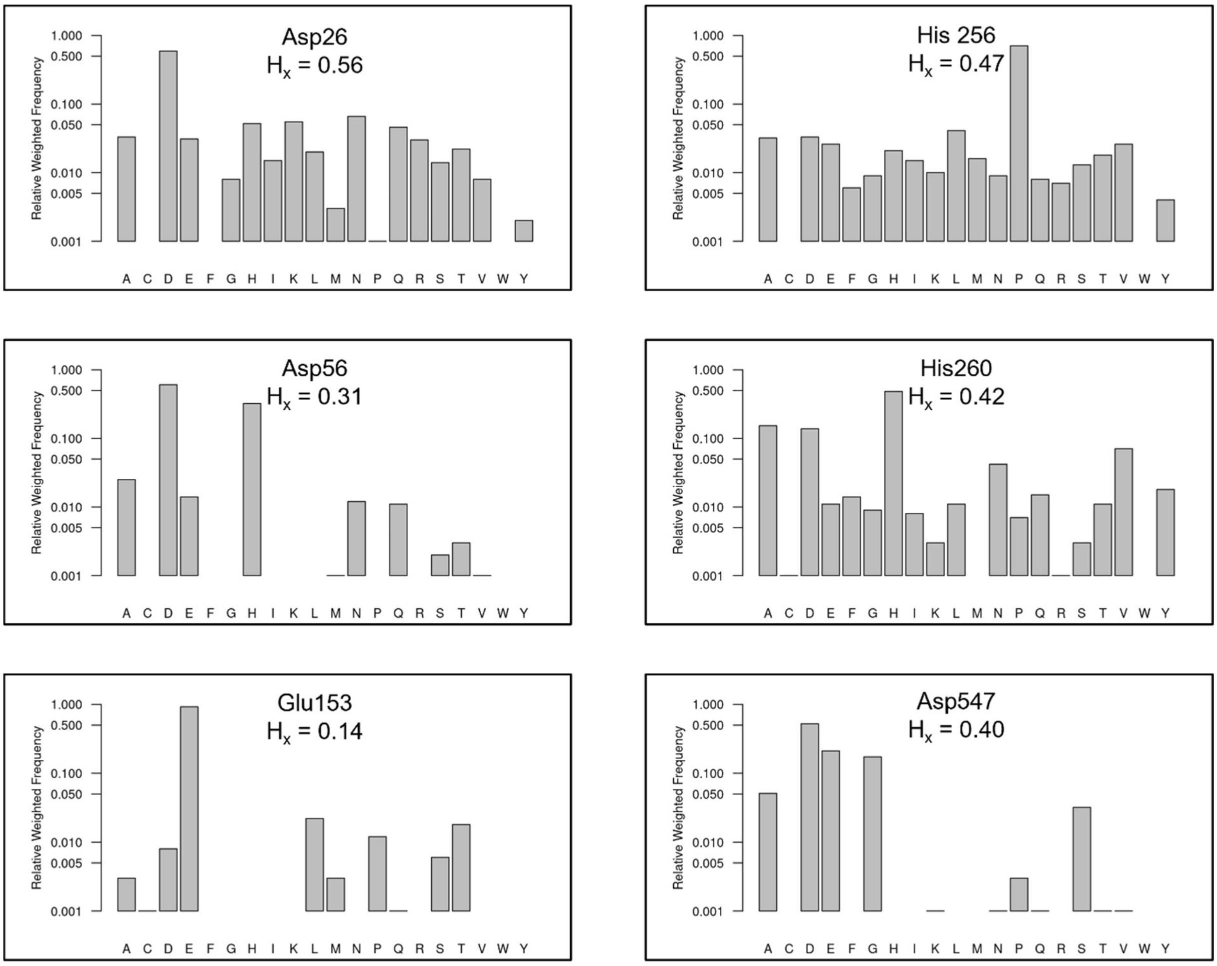


**Figure S6 MSA sequence-weighted residue frequency profiles at ionisable residue positions in Bad.F6Pkt.**

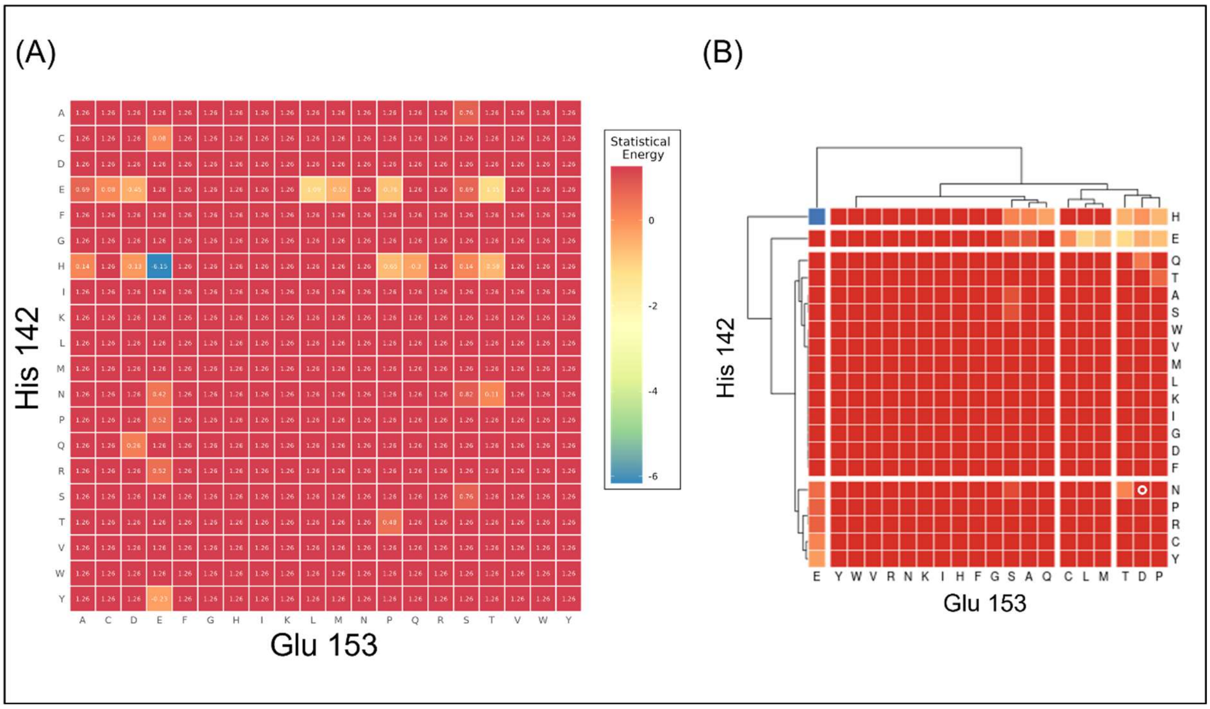


**Figure S7 Residue-pair statistical correlation energy heat maps for positions H142 and E153 in Bad.F6Pkt. (A)** Matrix heatmap annotated with statistical energies, calculated from sequence weighted observed residue type counts in a multiple sequence alignment as described in Methods using SEQUESTER [12]. **(B)** Hierarchical clustered heat map. The H142N:E153D double mutant is highlighted by the white circle.

#### Plasmids

**Table S5 Plasmids used in this study.**

| Plasmid | Short description | Source |
|---|---|---|
| pET-28a(+) | *ori f1*, $Kan^R$, T7 promoter | Novagen™ |
| pET28a_*Bad.f6pkt* | pET-28a(+) carrying *f6pkt* gene from *Bifidobacterium adolescentis* ATCC 15703 | [13] |
| pET28a_*Bad.f6pkt* H142N | pET-28a(+) carrying $f6pkt_{H142N}$ gene from *Bifidobacterium adolescentis* ATCC 15703 | [13] |
| pET28a_*Bad.f6pkt* H548N | pET-28a(+) carrying $f6pkt_{H548N}$ gene from *Bifidobacterium adolescentis* ATCC 15703 | [13] |
| pET28a_*Bad.f6pkt* H548Y | pET-28a(+) carrying $f6pkt_{H548Y}$ gene from *Bifidobacterium adolescentis* ATCC 15703 | [13] |
| pET28a_*Bad.f6pkt* N549D | pET-28a(+) carrying $f6pkt_{N549D}$ gene from *Bifidobacterium adolescentis* ATCC 15703 | [13] |
| pET28a_*Bad.f6pkt* N549C | pET-28a(+) carrying $f6pkt_{N549C}$ gene from *Bifidobacterium adolescentis* ATCC 15703 | [13] |
| pET28a_*Bad.f6pkt* N549S | pET-28a(+) carrying $f6pkt_{N549S}$ gene from *Bifidobacterium adolescentis* ATCC 15703 | [13] |
| pET28a_*Bad.f6pkt* N549V | pET-28a(+) carrying $f6pkt_{N549V}$ gene from *Bifidobacterium adolescentis* ATCC 15703 | [13] |
| pET28a_*Ec.fsaA* L107Y:A129G | pET-28a(+) carrying $fsaA_{L107Y:A129G}$ gene from *Escherichia coli* K-12 MG1655 | Lab stock |
| pET28a_*Ps.lrhi* | pET-28a(+) carrying *lrhi* gene from *Pseudomonas stutzerii* | [13] |
| pET28a_*Gg.der* | pET-28a(+) carrying *der* gene from *Gallus gallus* | [13] |
| pET28a_*Gs.ackA* | pET-28a(+) carrying *ackA* gene from *Geobacillus stearothermophilus* DSM 22 | [13] |
| pET28a_*Go.gox0313* | pET-28a(+) carrying the codon-optimized *gox0313* gene from *Gluconobacter oxydans* DSM 2003 | Lab stock |
| pET28a_*Ms.nox* | pET-28a(+) carrying the *nox* gene from *Methanobrevibacter smithii* ATCC 35061 | Lab stock |
| pET28a_*Cs.glpK* | pET-28a(+) carrying *glpK* gene from *Cellulomonas sp.* NT3060 | [13] |

### Additional experimental results

#### Beneficial Bad.F6Pkt variant H548Y identified from screening of selected active site positions

In order to identify an active site-variant with improved affinity for GA, we screened a total of 20 variants in bound TPP-F6P adduct contacting positions Q321, S541, H548 and K605 of Bad.F6Pkt previously found to improve activity on the non-phosphorylated substrate D-fructose[13]. The majority of these active-site variants manifested a significantly impaired GA activity (Figure S8). In addition to the H548N mutation described previously[13], only the H548Y variant exhibited a significant increase in activity of 21 ± 8 % compared to the wild type enzyme. Therefore, the kinetic parameters of the H548Y Bad.F6Pkt variant were analyzed on GA and ERU (Table 2). With a $K_M$ value of 43.4 ± 3.5 mM for the $C_2$ aldehyde and 26.9 ± 1.9 mM for the $C_4$ sugar, the H548Y mutation significantly enhanced the apparent affinity for both substrates in comparison to the wild-type and the H548N variant (no substrate saturation up to 300 mM of GA[13]). The $k_{cat}/K_M$ of the H548Y mutant was determined to be 14.8 ± 1.0 $M^{-1}$ $s^{-1}$ for GA. Compared to H548N, an increase of 66 % in catalytic efficiency for ERU was observed, representing a 2.7-fold increase in comparison to the wild-type enzyme (Table 2, Figure 5).

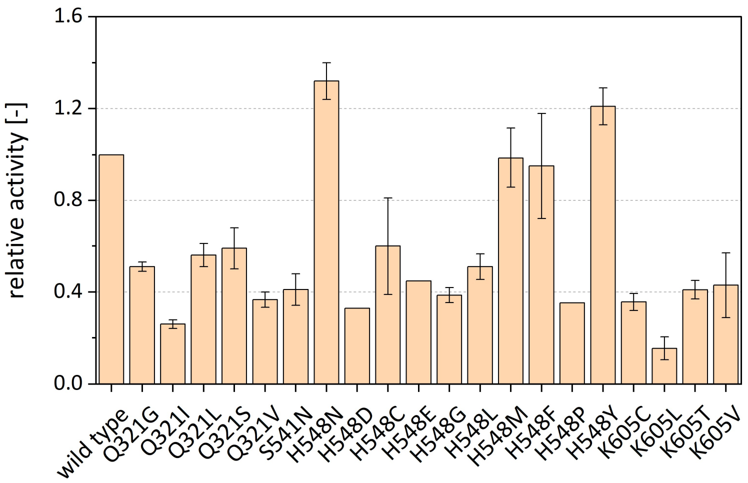


**Figure S8 Relative GA activity of Bad.F6Pkt active site single mutants.** The selection of mutations was based on their reported positive impact on activity towards the non-phosphorylated substrate D-fructose [13]. Beneficial variants, identified from screening of single mutation libraries, were produced in shake flasks, purified and their specific activity determined at pH 6.5, 37 °C in the presence of 200 mM GA. Relative activity is expressed as the ratio of the specific activities of the mutant and wild-type enzyme. Error bars indicate deviation of the mean (n ≥ 2).

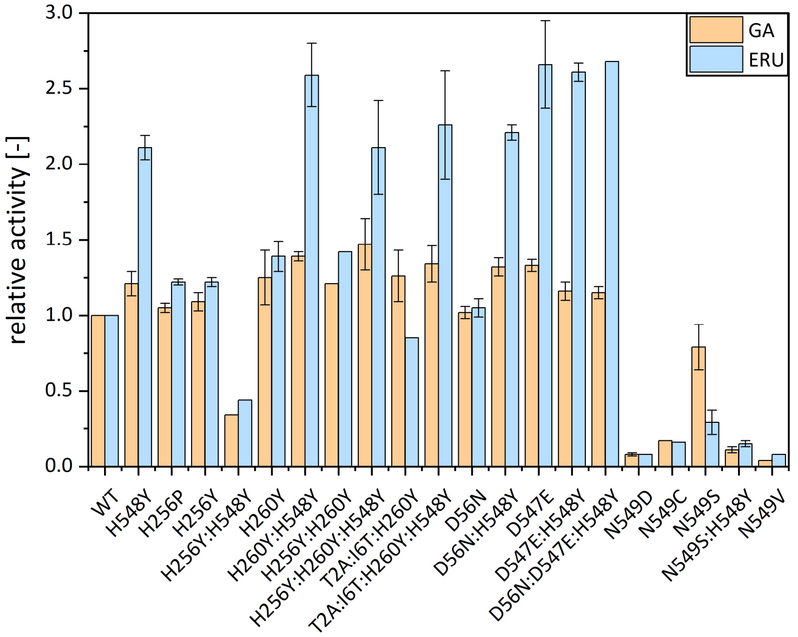


**Figure S9 Relative activity on D-erythrulose (ERU) and glycolaldehyde (GA) of Bad.F6Pkt variants carrying electrostatic mutations.** WT – wild type. Experiments were carried out at pH 6.5, 37 °C in the presence of 50 mM inorganic phosphate and 25 mM ERU or 200 mM GA. Relative activity is expressed as the ratio of the specific activities of the mutant and wild-type enzyme. Error bars indicate deviation of the mean (n ≥ 2).

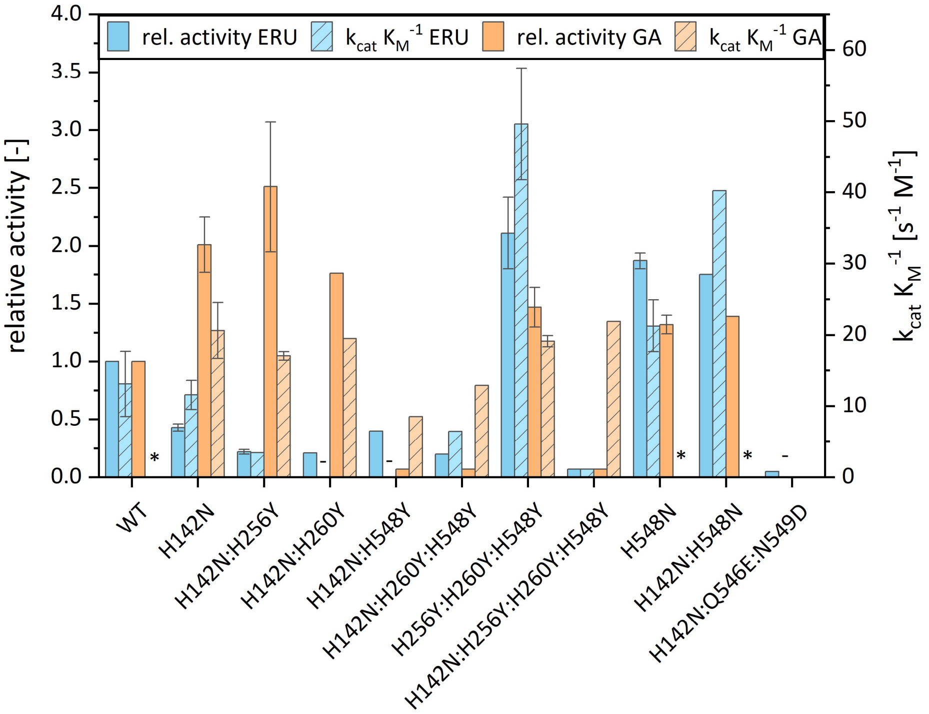


**Figure S10 Glycolaldehyde (GA) and erythrulose (ERU) activity and catalytic efficiency of Bad.F6Pkt H142N variants containing at least one other previously identified beneficial mutation for non-phosphorylated substrates .** Relative activity is expressed as the ratio of the specific activities of the mutant and wild-type enzyme at a substrate concentration of 25 mM ERU or 200 mM GA. Experiments were carried out at pH 6.5, 37 °C in the presence of 50 mM inorganic phosphate. Kinetic parameter fitting of experimental data to the Michaelis-Menten equation was carried out using the MATLAB R2023b curve fitting tool. In instances where substrate saturation was not observed under experimental conditions, $K_M$. and $k_{cat}$ /$K_M$ could not be determined (*). WT – wild type, (-) – not analysed. Error bars indicate deviation of the mean (n ≥ 2).

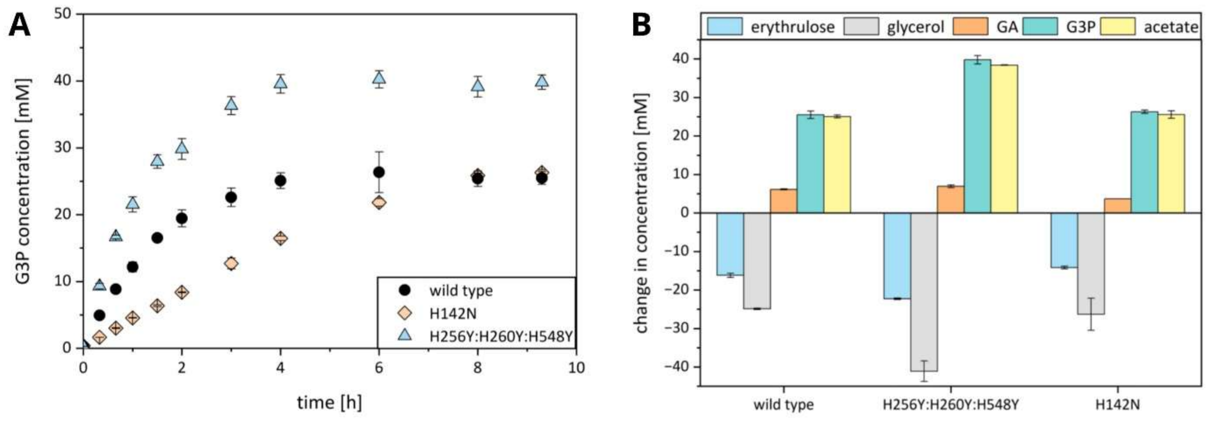


**Figure S11** ***In vitro*** **ATP regeneration from ERU by Bad.F6Pkt wild-type enzyme and different variants. A)** *In vitro sn*-glycerol 3-phosphate (G3P) production with ATP regenerated from ERU **B)** changes in substrate, intermediate and product concentrations observed between the start of the reaction and the final sample after 9.5 hours. A) + B) Experiments were carried out at pH 7.0, 37 °C in the presence of 50 mM sodium phosphate and glycerol, 0.8 mM TPP, 4 mM $MgCl_2$, 1 mM ADP and a starting concentration of 25 mM D-erythrulose. Phosphoketolase activity was regulated by employing either wild-type Bad.F6Pkt or indicated variants at a concentration of 2 mg $mL^{-1}$. Data represent mean and deviation of biological duplicates.

**Table S6 Inactivation of Bad.F6Pkt wild-type by D-threose and GA.** Phosphoketolase was incubated at pH 6.5 and 37 °C in the presence of 50 mM inorganic phosphate, 4 mM $MgCl_2$ and 1 mM TPP, as well as D-threose or GA at varying concentrations. Samples were taken at regular intervals and the remaining activity was immediately determined by the hydroxamate assay. Experimental data were fitted to a first-order inactivation model [14]. Irreversible enzyme inactivation by GA was investigated by J.-O. Kundoch (unpublished results). n ≥ 2

| substance | Concentration [mM] | Psuedo-First Order Rate Constant for Inactivation [$h^{-1}$] | Half-life [h] |
|---|---|---|---|
| D-threose | 10 | 0.036 ± 0.005 | 19.7 ± 2.7 |
| | 50 | 0.214 ± 0.023 | 3.3 ± 0.4 |
| glycolaldehyde | 10 | 0.415 ± 0.009 | 1.7 ± 0.0 |
| | 45 | 3.923 ± 0.118 | 0.2 ± 0.0 |
